# M6AFormer Prioritizes Unannotated Functional m^6^A Candidate Sites in the Human Epitranscriptome

**DOI:** 10.64898/2026.07.10.737679

**Authors:** Zhixin Niu, Chang Liu, Lei Gu

## Abstract

N^6^-methyladenosine (m^6^A) is a pervasive RNA modification with critical roles in post-transcriptional regulation, yet accurate transcriptome-wide identification of functional m^6^A sites remains challenging. Here, we present M6AFormer, a hybrid deep-learning framework that combines convolutional feature extraction with a lightweight Transformer to capture both local sequence motifs and broader contextual dependencies. M6AFormer consistently outperformed representative m^6^A predictors, including MST-M6A, CLSM6A and deepSRAMP. Transcriptome-wide scanning revealed a large repertoire of previously unannotated candidate m^6^A sites that retained hallmark m^6^A features, including canonical motif enrichment, characteristic spatial distribution and preferential overlap with m^6^A writer and reader binding regions. Importantly, M6AFormer-predicted sites were broadly associated with genetic and disease-relevant features, including SNPs, sequence variants and GWAS-linked loci, suggesting their potential contribution to human disease mechanisms. Finally, experimental validation confirmed a previously unreported m^6^A site in NEU4 mRNA and demonstrated its functional impact on cancer cell migration. Together, M6AFormer provides an accurate, interpretable and biologically informative framework for m^6^A site discovery.

## Introduction

N6-methyladenosine (m^6^A) is the most abundant internal modification in eukaryotic mRNA and represents a major layer of post-transcriptional gene regulation^1,2^. Its deposition, removal and recognition are mediated by methyltransferase complexes, demethylases and m6A-binding proteins, respectively^1,3^. Through these regulators, m^6^A influences multiple aspects of RNA metabolism, including pre-mRNA processing, nuclear export, translation, RNA stability and RNA-protein interactions^2,4–6^. Because these molecular effects are coupled to cell fate decisions, stress responses and disease-associated gene regulation, accurate identification of m^6^A sites is essential for understanding how transcriptome regulation is encoded and interpreted in different biological contexts.

Transcriptome-wide profiling has revealed several characteristic features of m^6^A methylation. m^6^A sites are frequently enriched in consensus DRACH sequence contexts and show non-random positional distributions along transcripts, particularly within coding regions, near stop codons and in untranslated regions^7,8^. However, the presence of a consensus motif alone is insufficient to determine methylation status^9^. Only a subset of adenosines within permissive sequence contexts are methylated, indicating that site selection depends on broader sequence grammar and transcript level features that may influence motif accessibility and recognition by regulatory factors^9–12^. This makes single nucleotide prioritization essential. M6AFormer addresses this need by learning discriminative sequence patterns from known m^6^A sites and applying them transcriptome wide to nominate candidate adenosines for downstream annotation, biological filtering and experimental validation.

Experimental technologies have provided indispensable maps of the m^6^A epitranscriptome, yet they remain limited for comprehensive site-level discovery^13,14^. Antibody-based MeRIP-seq/m^6^A-seq enabled transcriptome-wide profiling, but typically identify methylation-enriched regions rather than the precise modified nucleotide and can be affected by peak resolution, antibody dependence and changes in RNA abundance^14–16^. Crosslinking-based, chemical-conversion and enzyme-assisted approaches have improved single-nucleotide or quantitative mapping, but these assays can require specialized protocols, substantial sequencing depth, specific sample preparation or condition-matched experimental data^17,18^. More recently, nanopore direct RNA sequencing has enabled native RNA molecules to be interrogated without reverse transcription or amplification and can infer RNA modifications from raw current signals at the read or isoform level^19,20^. However, m^6^A detection from nanopore data remains dependent on computational models, sufficient transcript coverage, sequence context, control or training data, and caller-specific performance trade-offs^21^. As a result, experimentally validated m^6^A annotations remain incomplete across cell types, tissues, species and disease-relevant contexts. Computational prediction can help bridge this gap by prioritizing candidate sites for functional analysis, enabling transcriptome-scale screening and guiding experimental validation.

A number of computational approaches have been developed to predict m^6^A sites from RNA sequence and related features. Early predictors such as SRAMP used sequence-derived features and classical machine-learning models to identify mammalian m^6^A candidates^22^. More recent methods, including deep-learning frameworks such as CLSM6A, MST-A and deepSRAMP, have improved site-level prediction by learning richer representations of sequence, genomic position or cellular context^18,23,24^. These advances have made m^6^A prediction increasingly useful, but important challenges remain: many existing approaches are still centered on benchmark classification and do not fully bridge the gap between site prediction and biological discovery. In particular, predicted sites are rarely organized into a transcriptome-wide framework that distinguishes annotated from unannotated candidates, evaluates their regulatory and disease relevance, and prioritizes specific sites for experimental perturbation. A predictor intended for biological discovery should therefore be accurate and interpretable enough to support downstream mechanistic hypotheses.

To address these challenges, we developed M6AFormer, a CNN and lightweight Transformer-based framework for single-nucleotide-resolution prediction of m^6^A sites. M6AFormer was designed to capture both local sequence determinants and broader contextual features that may contribute to m^6^A site selection, while remaining suitable for transcriptome-scale application. In this study, we used M6AFormer to prioritize candidate m^6^A sites genome-wide and to connect model-predicted sites with canonical m^6^A features, regulatory protein occupancy, disease-associated genetic variation and targeted experimental validation. By integrating deep learning-based prediction with biological annotation and functional follow-up, this work aims to provide a practical framework for discovering candidate regulatory m^6^A sites and generating testable hypotheses in RNA biology and disease.

## Results

### M6AFormer was designed to reveal hidden functional m^6^A sites beyond current annotations

We developed M6AFormer to address a central limitation in m^6^A biology: current experimental maps and annotation resources capture only a fraction of the potentially functional m^6^A landscape. Many regulatory m^6^A sites may remain underrepresented because profiling experiments are condition-dependent, limited by sequencing depth, affected by antibody or protocol biases, or absent from the cell states currently represented in public databases^15,25^. We therefore developed M6AFormer as an intelligent discovery framework for nominating candidate m^6^A regulatory sites that are missed or insufficiently represented by existing annotations, while further linking these predictions to regulatory context, disease relevance, and experimental validation. (Fig. 1).

**Fig. 1:**
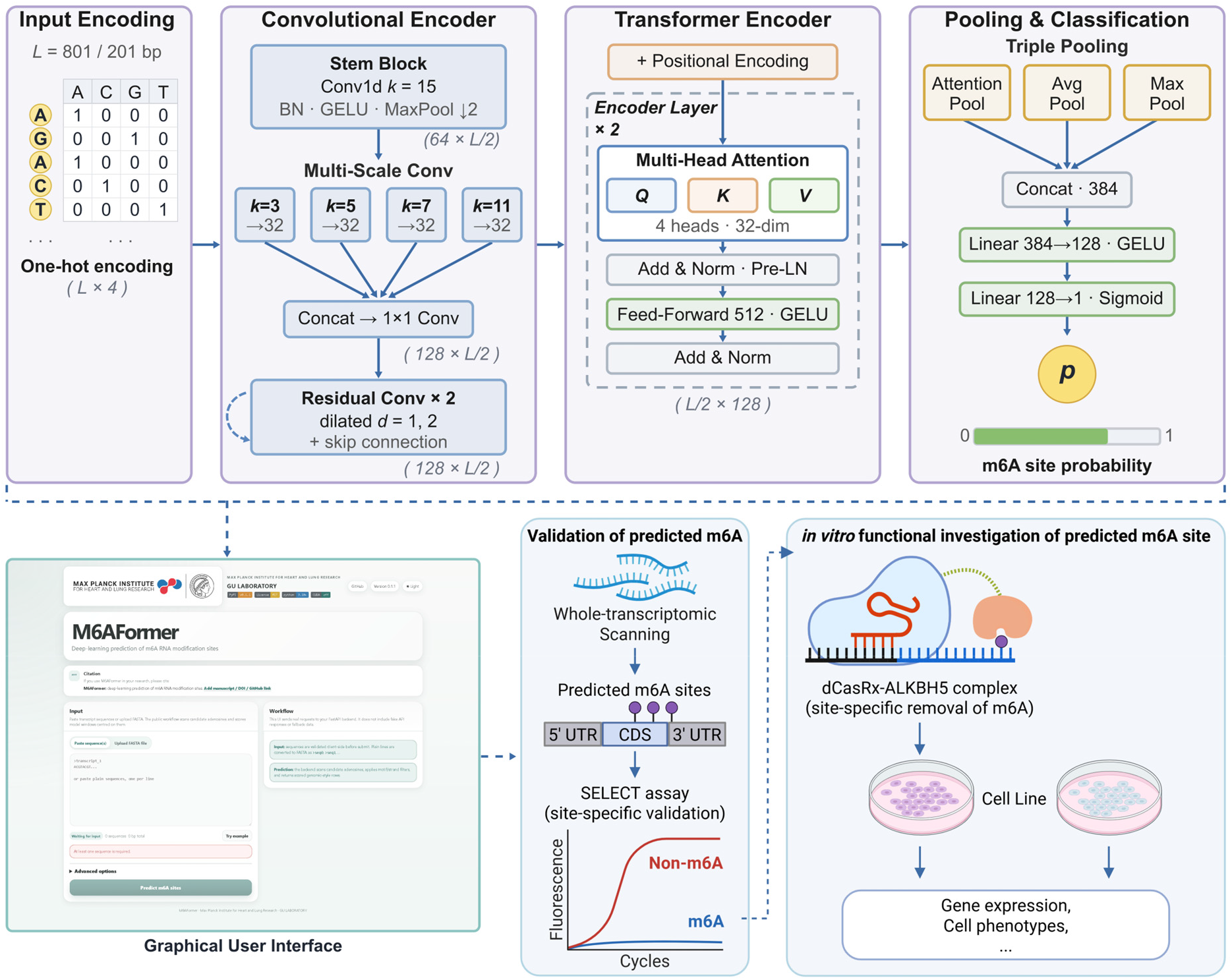
Overview of the M6AFormer architecture and analysis workflow. Schematic overview of M6AFormer and downstream analyses. Candidate adenosine-centered RNA sequences are one-hot encoded and processed through a convolutional encoder, transformer encoder and triple-pooling classifier to generate a site-level m^6^A probability. The lower workflow shows the GUI for sequence submission and prediction, transcriptome-wide scanning of candidate m^6^A sites across transcript regions, SELECT-based site validation and functional assays after dCasRx-ALKBH5-mediated m^6^A removal. SELECT, single-base elongation- and ligation-based qPCR amplification method

M6AFormer takes adenosine-centered nucleotide windows as input and produces site-level m^6^A probabilities through a CNN-transformer architecture that combines motif-scale convolution, multi-scale contextual encoding, self-attention and pooled sequence representations (Fig. 1). This architecture enables the model to capture local m^6^A sequence grammar while also considering broader sequence context that may not be represented by a single canonical motif. To make the predictions usable for biological discovery, we implemented a user-friendly graphical interface (GUI) and linked model scores to transcriptome-wide scanning, database annotation, functional and disease enrichment, SELECT-based m^6^A validation and targeted dCasRx-ALKBH5 perturbation. Thus, the framework converts sequence-level evidence into experimentally tractable hypotheses about hidden m^6^A regulatory sites.

### Sequence modeling provides a reliable basis for candidate prioritization

We first evaluated whether M6AFormer robust enough to support downstream discovery. CLSM6A, MST-m6A, and deepSRAMP were selected as representative m^6^A predictors because each model was accompanied by clearly defined datasets, accessible source code, and released pretrained weights. M6AFormer was then benchmarked against these predictors using harmonized evaluation metrics across receiver operating characteristic curves, precision-recall curves, and multi-metric radar summaries (Supplementary Table S1). We evaluated both all-adenosine and DRACH-restricted settings using 201-bp and 801-bp input windows. These four benchmark configurations are denoted as ALL201 and ALL801 for the all-adenosine tasks, and DRACH201 and DRACH801 for the DRACH-restricted tasks.

M6AFormer achieved strong performance across these settings (Fig. 2A-B; Supplementary Table S2-3). In the ALL801 configuration, the model achieved an AUROC of 0.984 and an AUPRC of 0.980, with an accuracy of 0.949, F1 score of 0.950 and Matthews correlation coefficient (MCC) of 0.898. The ALL201 model also performed well, reaching an AUROC of 0.978 and an AUPRC of 0.972. As expected, performance decreased in the more stringent DRACH-restricted setting but remained robust. The DRACH801 model achieved an AUROC of 0.917 and an AUPRC of 0.916, with an MCC of 0.666. The DRACH201 model reached an AUROC of 0.856 and an AUPRC of 0.861. These results indicate that broader sequence context improves candidate prioritization and that M6AFormer retains discriminative information even when negatives are restricted to motif-compatible adenosines. Compared with the evaluated predictors under the matched DRACH-restricted setting, M6AFormer produced stronger ROC and precision-recall profiles (Fig. 2C, D). This difference was statistically significant rather than a product of sampling variation. By the DeLong test for two correlated ROC curves, M6AFormer in the DRACH-restricted 801-nt configuration reached an AUROC of 0.917, compared with 0.769 for the CNN baseline, 0.745 for CLSM6A, 0.599 for MST-m6A and 0.563 for deepSRAMP, and all pairwise differences were significant by a two-sided test (P < 2.2 × 10⁻¹^6^; Supplementary Table S4). Each comparator reproduced its own published performance on its own held-out data (Supplementary Table S5), so the margin observed here reflects cross-dataset generalization rather than an incorrect re-implementation. The precision-recall improvement is particularly relevant for discovery of unannotated candidate sites, because experimental follow-up usually begins from a limited number of high-confidence predictions. Radar plots further showed that M6AFormer maintained balanced performance across AUROC, average precision, MCC, accuracy, sensitivity and specificity (Fig. 2E).

**Fig. 2:**
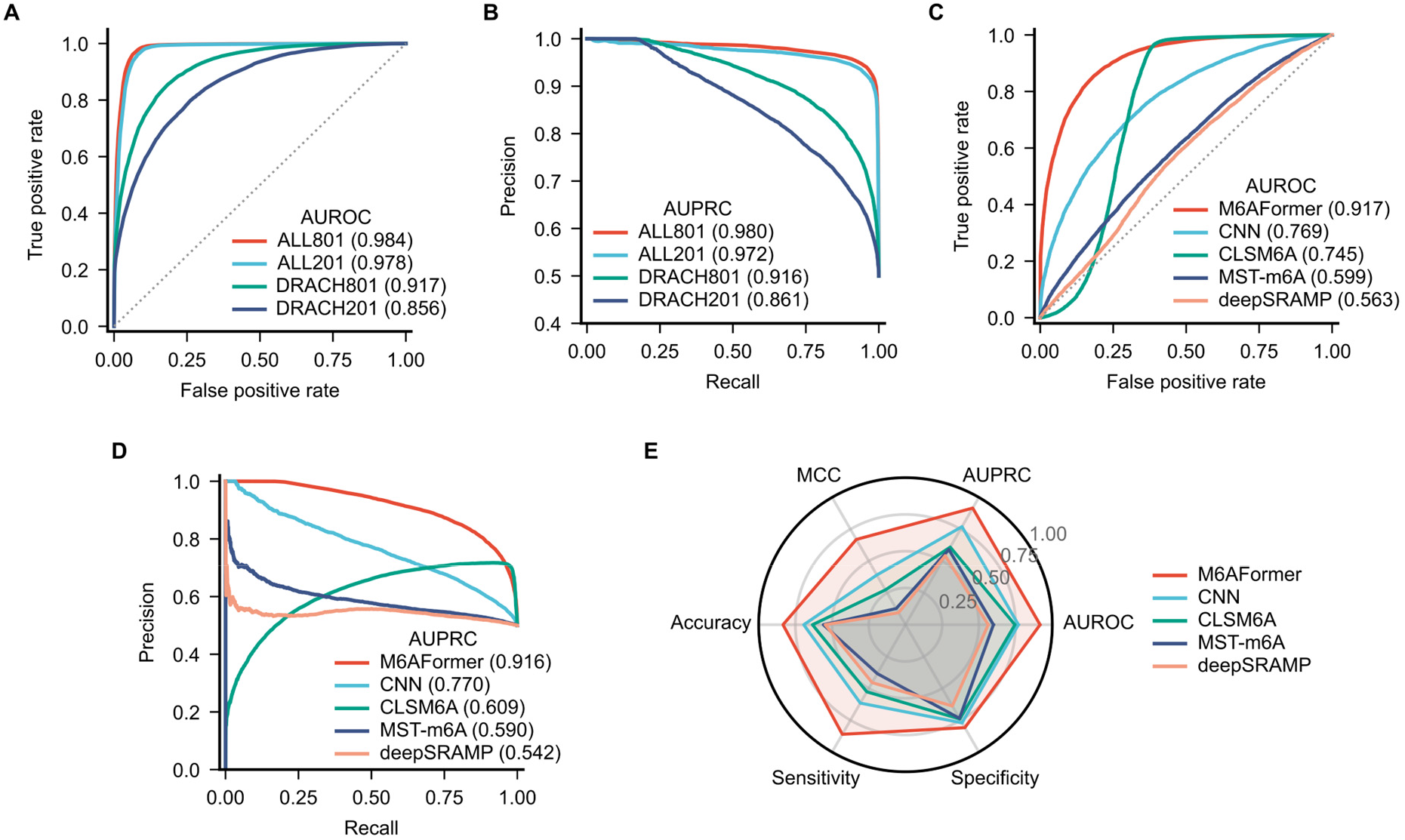
Benchmark comparison of M6AFormer and existing m6A prediction models. **(A)** Receiver operating characteristic and **(B)** precision-recall curves for the four M6AFormer configurations (ALL201, ALL801, DRACH201, DRACH801) on the held-out test set (n = 27,782). **(C)** Receiver operating characteristic and **(D)** precision-recall curves for M6AFormer (DRACH-801) against a CNN baseline, CLSM6A, MST-m6A and deepSRAMP on the same held-out sites, each method scoring every site from sequence alone. Curves show true positive rate versus false positive rate **(A, C)** or precision as a function of recall **(B, D)**; values in parentheses indicate AUROC or AUPRC. **(E)** Radar plot summarizing AUROC, AUROC, MCC, accuracy, sensitivity and specificity for the same cross-method comparison. Colors denote models or configurations as indicated in the panel legends.

We further compared these predictors under matched cell-type conditions, evaluating each competitor with its own cell-line-specific model on the corresponding cell-line subset of the held-out data, alongside M6AFormer and the CNN baseline (Supplementary Fig. S1; Supplementary Table S6). M6AFormer generalized consistently across cell lines, with an AUROC of 0.93 to 0.95 and an MCC near 0.73 across A549, HEK293T, HeLa and HepG2. This exceeded the CNN baseline and the dedicated cell-line-specific models of MST-m6A, CLSM6A and deepSRAMP, several of which approached random discrimination on individual cell lines. A single general M6AFormer model therefore outperformed competitors even when those competitors were given cell-line-matched models.

### Ablation and attribution analyses support biologically plausible sequence learning

We next examined whether the performance of M6AFormer depended on the intended architecture and whether its predictions were supported by interpretable sequence evidence. Ablation analysis, performed in the DRACH-restricted 801-nt configuration, showed that each major component contributed to performance (Fig. 3A-C, Supplementary Table S7). In the ablation benchmark, the full model achieved an AUROC of 0.916, an AUPRC of 0.916, an MCC of 0.662 and an accuracy of 0.829. Removing individual components reduced performance to different extents. The strongest decrease occurred when the transformer encoder was removed. Without the transformer, AUROC decreased to 0.857 and AUPRC decreased to 0.832, with MCC falling to 0.566. This result suggests that self-attention contributes information beyond local motif recognition. Removing the residual convolutional module also reduced performance, with AUROC decreasing to 0.904 and MCC to 0.635, indicating that this component contributes to overall performance. Removing multi-scale convolution caused a smaller but consistent decrease, with AUROC of 0.910 and MCC of 0.648. Removing attention pooling similarly reduced performance, with AUROC of 0.910 and MCC of 0.653. Thus, the full architecture improved prioritization by combining motif-scale recognition, expanded convolutional context, transformer-based sequence modeling and adaptive sequence summarization.

**Fig. 3:**
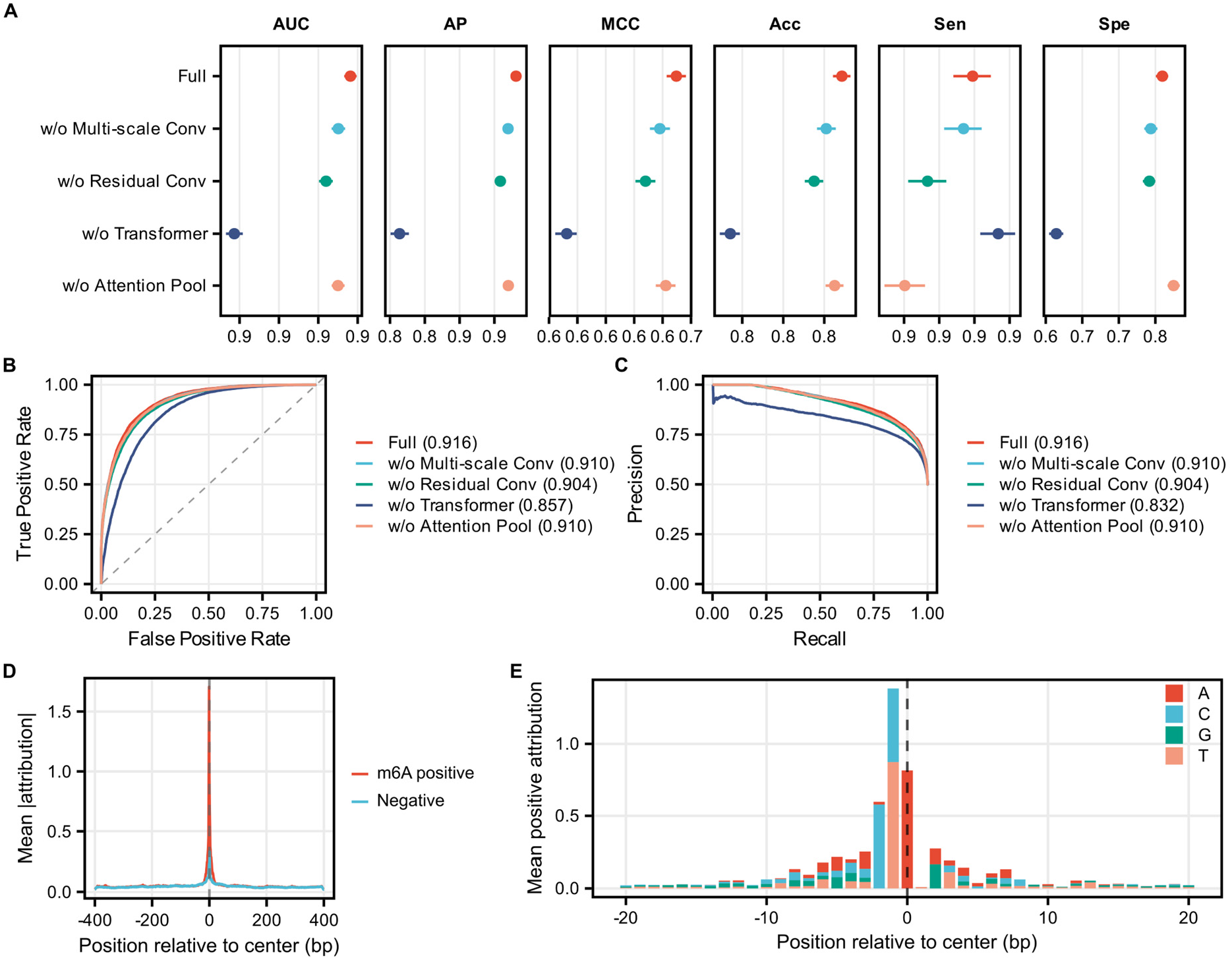
Ablation and attribution analysis of M6AFormer. **(A)**. Ablation analysis comparing the full M6AFormer model with variants lacking multi-scale convolution, residual convolution, the transformer encoder or attention pooling. Points show performance metrics; horizontal bars indicate bootstrap confidence intervals. **(B)**. Receiver operating characteristic curves for the full and ablated models; values in parentheses indicate AUROC. **(C)**. Precision-recall curves for the same ablation models; values in parentheses indicate AUPRC. **(D)**. Position-wise attribution profile across the input sequence for m6A-positive and negative sites. The dashed vertical line marks the candidate adenosine. **(E)**. Mean positive nucleotide attribution around the candidate site, separated by nucleotide identity. Bars show attribution contribution by base.

Attribution analysis further indicated that model decisions were anchored in biologically plausible sequence features rather than diffuse background signal. Position-wise attribution showed a pronounced importance peak centered on the candidate adenosine and the immediately surrounding nucleotides (Fig. 3D). Nucleotide-level attribution highlighted base-specific contributions around the central site (Fig. 3E). These patterns are consistent with established m^6^A sequence grammar while allowing additional contextual information to influence the final score. For unannotated-site discovery, this provides a useful safeguard because high-scoring sites can be assessed for plausible sequence support before experimental follow-up.

Together, these results indicate that M6AFormer predictions are anchored in sequence features consistent with established m^6^A grammar, and that each architectural component contributes measurably to predictive performance.

### Transcriptome-wide scanning nominates a large unannotated m^6^A-like candidate landscape

We next applied M6AFormer to transcriptome-wide discovery to identify candidate m^6^A sites not fully represented in existing annotations. Because a genome-wide scan scores every adenosine rather than only motif-restricted sites, we used the all-adenosine 801-nt model (ALL801), whose training negatives span the full genomic adenosine population. Predicted sites were annotated against existing m^6^A resources, including RMBase and m6A-Atlas, and classified by database annotation status, DRACH motif status and transcript region. An intersection analysis of the full predicted set showed that database-supported predictions were largely shared between RMBase and m6A-Atlas, whereas the majority of predicted adenosines fell outside both catalogs (Supplementary Fig. S2). The transcriptome-wide scan recovered both database-supported (annotated) and predicted candidate (unannotated) sites by discriminating each adenosine on its predicted probability against the best-F1 operating threshold of the model (ALL801, probability = 0.48).

To define a high-confidence subset of predicted m6A sites for downstream analyses, we compared candidate probability cutoffs (prob > 0.9, 0.95, 0.99, and 0.995) with respect to the model predictive precision, probability calibration and site yield on the independent held-out test set (Supplementary Fig. S3A-C). Precision (positive predictive value, PPV) increased monotonically with the cutoff and saturated at prob > 0.99 (PPV = 0.992, corresponding to a 0.8% FDR), a value reproduced in the external validation set, and predicted probabilities were well calibrated across the full range (Supplementary Fig. S3B, C). Raising the cutoff further to 0.995 improved precision by only 0.03 (PPV = 0.995) while discarding ∼4.4-fold more sites (80,497 to 18,465 retained), whereas looser cutoffs 0.9/0.95 admitted substantially more false positives (PPV = 0.936/0.962; Supplementary Fig. S3A, B). These results showed that prob > 0.99 offered the best balance between predictive precision and site yield to define high-confidence m^6^A sites. At the high-confidence threshold, an intersection analysis showed the same overlap structure seen for the full set, with database-supported predictions largely shared between RMBase and m6A-Atlas and, at this stricter cutoff, the unannotated candidates forming a distinct group outside both catalogs (Fig. 4A). The high-confidence set comprised 312,625 sites, of which 74.3% (232,128) were database-annotated and 25.7% (80,497) were unannotated (Fig. 4B). Thus, M6AFormer predominantly recovers known m^6^A sites, while still nominating 80,497 candidates absent from current m^6^A databases.

**Fig. 4:**
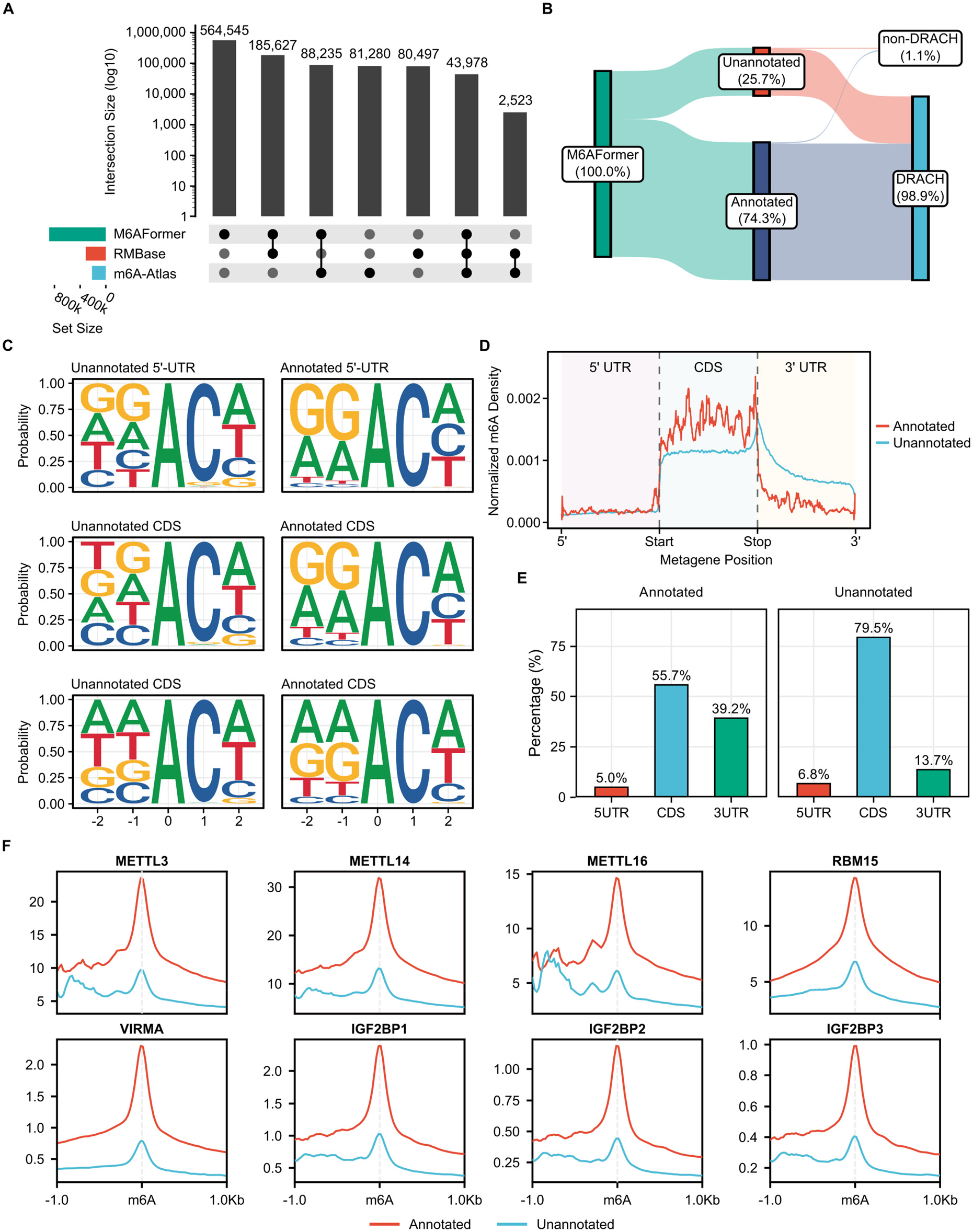
Transcriptome-wide annotation of M6AFormer-predicted m6A sites. **(A)**. UpSet-style overlap analysis comparing high-confidence M6AFormer-predicted sites (probability ≥ 0.99) with RMBase and m6A-Atlas. Bar heights show intersection sizes (log10), filled dots mark the set combination represented by each bar, and set sizes are shown at left.**(B)**. Sankey diagram summarizing the database annotation status and DRACH motif status of high-confidence predicted sites. Percentages show the fraction of sites in each category. **(C)**. Sequence logos for annotated and unannotated predicted sites across transcript regions. Letter height shows nucleotide frequency at each position relative to the candidate adenosine. **(D)**. Metagene distribution of annotated and unannotated predicted sites across 5’ UTR, coding sequence and 3’ UTR. **(E)**. Transcript-region distribution of annotated and unannotated predicted sites. Bar labels show percentages. **(F)**. m6A writer, regulator and reader signal around predicted m6A sites. Red and blue lines represent annotated and unannotated sites, respectively. The dashed vertical line marks the predicted m6A site.

We next asked whether these predictions retained m^6^A-like sequence properties. Among high-confidence, 98.9% contained a DRACH motif and only 1.1% did not (Fig. 4B); the unannotated high-confidence candidates were likewise almost entirely DRACH-containing (98.1%). Motif logos showed recognizable m^6^A-associated sequence patterns across transcript regions in both annotated and unannotated sets, although their motif composition differed (Fig. 4C). These observations indicate that many unannotated predictions are not arbitrary adenosines. Rather, they include candidate sites with sequence features consistent with known m^6^A biology, along with additional non-canonical or context-dependent candidates that may be underrepresented in current resources. Transcript position analysis further supported the biological plausibility of model-nominated sites. Annotated predictions showed the expected enrichment within coding sequence and around transcript landmarks, including a prominent metagene pattern near the coding region and stop codon (Fig. 4D). Unannotated predictions displayed a distinct but structured distribution, with a greater proportion in coding sequence. Region annotation showed that annotated predictions were distributed across 5’ UTR, CDS and 3’ UTR regions at 5.0%, 55.7% and 39.2%, respectively, whereas unannotated predictions were distributed at 6.8%, 79.5% and 13.7% (Fig. 4E).

To connect unannotated candidate sites with m^6^A regulatory machinery, we examined public binding or profiling signals for m^6^A writers, associated regulators and readers around predicted sites. Annotated sites showed strong enrichment around METTL3, METTL14, METTL16, RBM15, VIRMA and IGF2BP family proteins, as expected for bona fide m^6^A-associated regions (Fig. 4F). Importantly, unannotated sites also showed local signal enrichment, although generally weaker than annotated sites. This supports the interpretation that a subset of model-nominated unannotated sites are positioned within m^6^A regulatory contexts. The enrichment of unannotated predictions near m^6^A machinery signals supports the second interpretation for a subset of sites and provides a rationale for carrying these *candidates into functional and disease-oriented analyses*.

### Unannotated candidate sites are linked to functional programs and disease-associated variation

We next investigated whether M6AFormer-nominated sites, particularly the unannotated candidate set, were enriched for functional and disease-associated programs. Both annotated and unannotated predicted-site gene sets were enriched for biologically relevant pathways. KEGG enrichment included ribosome-related pathways, neurodegeneration, neuroactive ligand signaling, lysosome biogenesis, endocytosis, calcium signaling, cadherin signaling and axon guidance (Fig. 5A). GO Biological Process enrichment highlighted RNP complex biogenesis, mitotic cell cycle regulation, Wnt signaling, ribosome biogenesis, mitochondrial organization, cell adhesion, synaptic transmission and axonogenesis (Fig. 5B). The presence of RNA-processing, translational, neuronal, cell-cycle and adhesion-related terms suggests that M6AFormer-nominated sites mark transcripts involved in regulatory programs where m^6^A has plausible functional consequences.

**Fig. 5:**
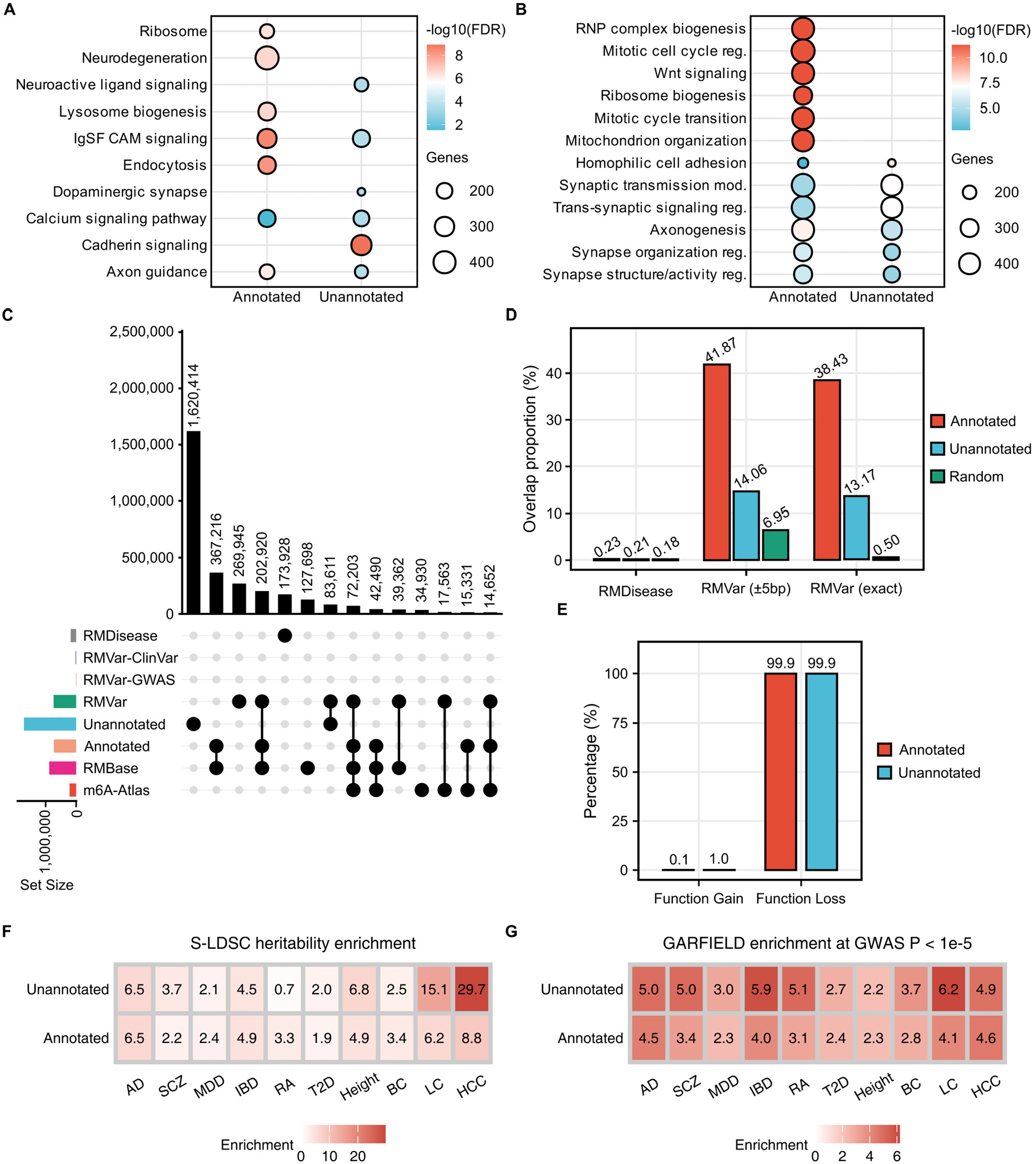
Functional enrichment and disease-variant annotation of predicted m6A sites. **(A)**. KEGG pathway enrichment of genes associated with annotated and unannotated M6AFormer-predicted sites. Dot size shows the number of genes in each term, and color shows enrichment strength as shown in the legend. **(B)**. GO Biological Process enrichment for the same gene sets. Terms are grouped by annotated and unannotated predicted-site categories. **(C)**. Overlap analysis integrating M6AFormer predictions with RMBase, m6A-Atlas, RMVar and RMDisease resources. Bar heights show intersection sizes; filled dots mark the resource combination. **(D)**. Overlap proportion between annotated, unannotated and random sites with RMDisease and RMVar annotations. RMVar overlaps are shown for exact matches and +/-5 bp windows. **(E)**. Predicted functional consequence distribution of annotated and unannotated sites based on RMVar annotations. **(F)**. S-LDSC regression heritability enrichment for annotated and unannotated predicted-site annotations across disease and complex traits. **(G)**. GARFIELD enrichment of annotated and unannotated annotations at GWAS P < 1e-5.

Disease-focused enrichment analyses gave a concordant view of the candidate-site gene sets. DO enrichment of annotated-site genes included intellectual disability and multiple cancer-related categories, including liver carcinoma, hepatocellular carcinoma, respiratory system cancer, and lung cancer (Supplementary Fig. S4A). For unannotated-site genes, DO terms were enriched for neurodevelopmental and neurological categories, including pervasive developmental disorder, autism spectrum disorder, focal epilepsy and temporal lobe epilepsy, together with reproductive and sensory categories such as spermatogenic failure, male infertility and nonsyndromic deafness (Supplementary Fig. S4A). DGN enrichment similarly highlighted neurodevelopmental, anthropometric, respiratory and behavioral traits, including autistic behavior, waist-hip ratio, vital capacity, smoking-related traits, scoliosis-related terms, narcolepsy and seizure-related categories (Supplementary Fig. S4B). These results indicate that the unannotated candidate set captures transcripts involved in organized biological and disease-associated programs rather than an unstructured collection of sequence matches.

We then examined whether model-nominated sites overlap disease-associated RNA modification resources. M6AFormer predictions were compared with RMDisease and RMVar annotations, including exact RMVar overlaps and overlaps within 5 bp (Fig. 5C, D). Annotated predicted sites showed strong overlap with RMVar, as expected from their database support. At the 0.99 threshold, 13.17% of unannotated sites overlapped RMVar exactly and 14.06% overlapped RMVar within 5 bp, compared with 0.50% and 6.95% for random background, respectively, indicating that the disease-resource overlap is not confined to already annotated m^6^A sites, but extends to model-nominated candidates absent from current m^6^A annotations. Both annotated and unannotated candidate sets were dominated by predicted function-loss annotations in RMVar-derived categories (Fig. 5E). These annotations do not establish the function of every individual site, but they indicate that M6AFormer-prioritized regions overlap positions where RNA modification changes may have regulatory consequences.

To evaluate whether predicted m^6^A sites are enriched for common disease genetic signals, we performed stratified LD-score regression (S-LDSC) and GARFIELD analyses across multiple traits (Fig. 5F, G; Supplementary Table S8-9). Stratified LD-score regression revealed heritability enrichment across several disease and complex-trait GWAS, including Alzheimer’s disease (AD), schizophrenia (SCZ), inflammatory bowel disease (IBD), height, lung cancer (LC) and hepatocellular carcinoma (HCC). In several traits, unannotated M6AFormer sites showed enrichment comparable to or greater than annotated sites including LC and HCC where unannotated predictions reached enrichments of 15.1 (LC) and 29.7 (HCC), exceeding the corresponding annotated estimates (6.2 and 8.8) (Fig. 5F; Supplementary Table S8). GARFIELD analysis provided an orthogonal enrichment framework based on GWAS association thresholds and local genomic covariate adjustment. The direction of enrichment was broadly consistent with S-LDSC results at an association threshold of P < 1 × 10^-5^, strengthening the evidence that M6AFormer-nominated sites capture disease-relevant regulatory variation (Fig. 5G; Supplementary Table S9). Together with pathway enrichment and RMVar/RMDisease overlap, these analyses support the view that the unannotated candidate landscape is enriched for disease-relevant regulatory information. They do not identify causal variants or establish disease mechanisms at individual loci; rather, they provide convergent prioritization evidence that can guide mechanistic follow-up.

### Experimental validation confirms model-nominated sites and functionally perturbs a prioritized m^6^A regulatory site

Finally, we tested whether M6AFormer-nominated candidates could be detected experimentally and whether a prioritized site could be functionally interrogated. We first performed SELECT assays across multiple model-nominated candidate sites (Fig. 6A). For each candidate, relative m^6^A abundance was calculated by comparing the predicted m^6^A site with an experimentally supported non-methylated adenosine site within the same transcript, providing a transcript-matched negative control for site-level detection. Several M6AFormer-nominated sites showed increased relative m^6^A abundance over their matched control adenosines, supporting the ability of the framework to nominate experimentally measurable m^6^A candidates.

**Fig. 6:**
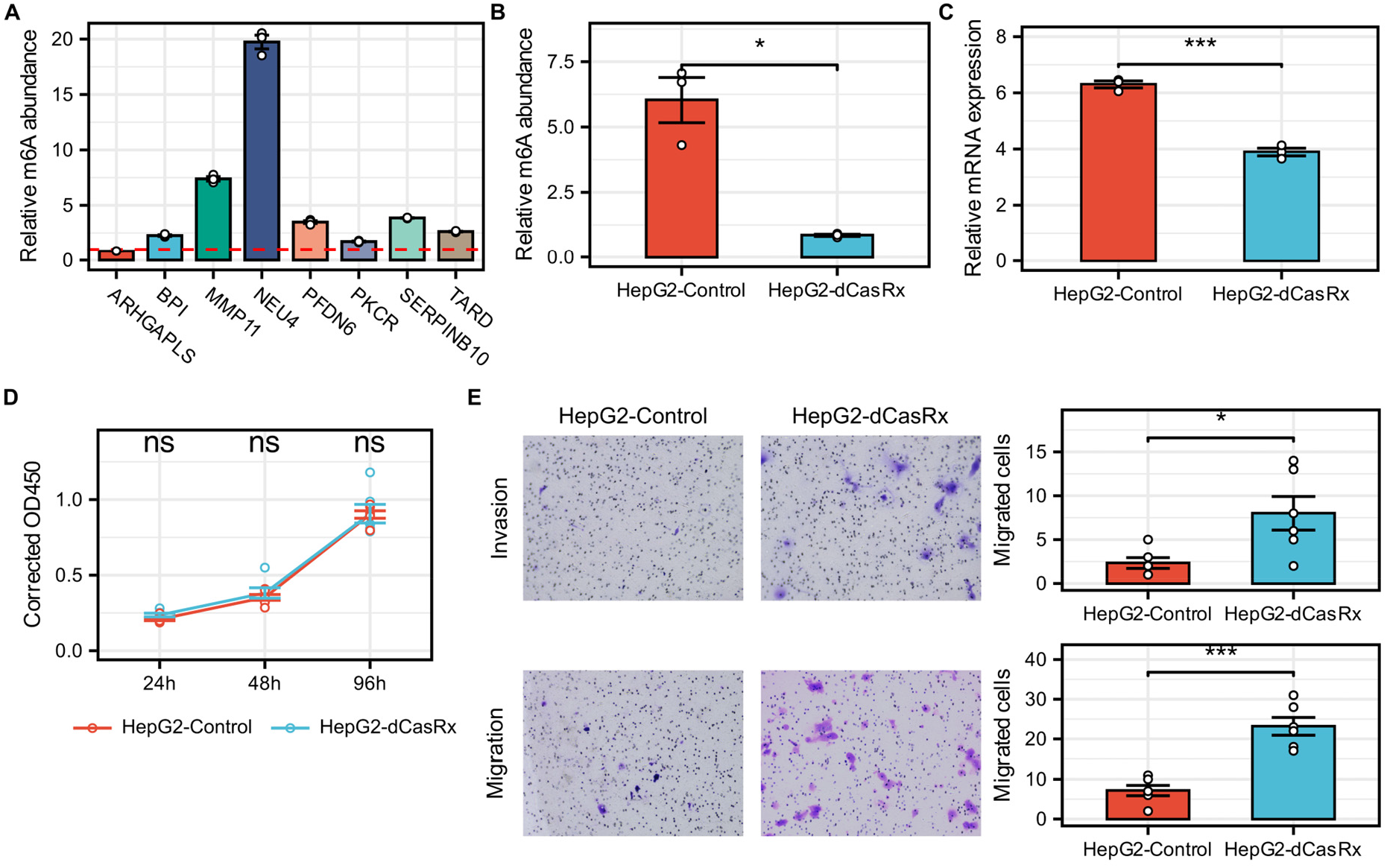
SELECT validation and dCasRx-ALKBH5 perturbation of prioritized m6A sites. **(A)**. SELECT validation of candidate m6A sites prioritized by M6AFormer. Bars show relative m6A abundance for the indicated candidate genes; the dashed red line marks the reference threshold used for comparison. Points denote experimental replicates. **(B)**. SELECT measurement of NEU4 m6A abundance after dCasRx-ALKBH5 targeting in HepG2 cells. **(C)**. NEU4 mRNA expression measured by qPCR in control and dCasRx-ALKBH5-treated HepG2 cells. **(D)**. CCK8 proliferation assay comparing control and dCasRx-ALKBH5-treated cells over time. Points denote replicates; lines connect group means. **(E)**. Representative transwell invasion and migration images and quantification. Bars show migrated or invaded cell counts, points denote replicate fields or biological replicates, and error bars indicate s.e.m. Statistical annotations indicate ns, not significant; *P < 0.05; ***P < 0.001.

For targeted perturbation, we focused on a high-confidence candidate site in NEU4 in HepG2 cells. NEU4 was well expressed in HepG2 in the cell-line expression matrix, providing sufficient transcript abundance for site-level SELECT detection (Supplementary Fig. S5). This target also matched the disease context emerging from the transcriptome-wide prioritization analyses, in which unannotated M6AFormer sites showed one of the strongest stratified LD-score enrichment signals for HCC (enrichment = 29.7, compared with 8.8 for annotated sites; Fig. 5F). Beyond these prioritization criteria, NEU4 has documented clinical relevance in liver cancer, acting as a tumour suppressor that restrains HCC cell motility and metastasis, being markedly downregulated in HCC tissues, and showing an association between reduced expression and both advanced disease and poorer patient survival (Supplementary Fig. S6) ^26^. The NEU4 candidate therefore combined high model confidence, detectable expression in the corresponding cell line, HCC-relevant prioritization evidence and established clinical significance in HCC, making it a suitable target for dCasRx-ALKBH5 perturbation.

We used a CRISPR-dCasRx-ALKBH5 system to remove m^6^A in a site-directed manner at the prioritized NEU4 region. SELECT analysis confirmed that dCasRx-ALKBH5 perturbation substantially reduced local m^6^A abundance compared with control cells (Fig. 6B), demonstrating that the candidate site could be experimentally edited. We next measured NEU4 mRNA expression and found that dCasRx-ALKBH5-mediated m^6^A removal decreased NEU4 transcript abundance (Fig. 6C). This result suggests that methylation at or near the prioritized site contributes to NEU4 RNA regulation in HepG2 cells. We then asked whether targeted perturbation of this prioritized m^6^A site was associated with cellular phenotypes. CCK8 assays showed no significant difference in proliferation between control and dCasRx-ALKBH5-treated cells across the measured time points (Fig. 6D), consistent with the reported role of NEU4 as a regulator of cell adhesion and motility rather than a primary driver of proliferative growth. In transwell assays, dCasRx-ALKBH5 perturbation increased both invasion and migration compared with control cells (Fig. 6E). These phenotypic changes indicate that removal of m^6^A at the prioritized NEU4 site is associated with altered tumor-relevant cellular behaviour.

Together, the experimental results close the discovery loop established by M6AFormer. Multi-site SELECT validation showed that model-nominated candidates can correspond to detectable m^6^A sites. Targeted dCasRx-ALKBH5 perturbation then demonstrated that a prioritized site can be edited, linked to transcript abundance and associated with cell migration and invasion phenotypes. We do not infer from these data that the NEU4 site alone fully explains the observed phenotypic changes. Rather, the results provide direct evidence that M6AFormer can reveal unannotated, functionally relevant m^6^A regulatory sites suitable for mechanistic follow-up.

## Discussion

M6AFormer was developed to address the incompleteness of current m^6^A annotations and to prioritize candidate regulatory sites that remain underrepresented in existing experimental maps. Our central observation is that pairing sequence-based prediction with transcriptome-wide annotation, disease-variant enrichment and targeted validation can prioritize candidate m^6^A sites that lie beyond current annotations but are supported by convergent biological and genetic evidence. Rather than functioning only as another classifier, M6AFormer learns sequence features consistent with known m^6^A grammar, nominates candidate sites across the transcriptome, and links a prioritized site to methylation abundance, transcript expression and cellular phenotype. The framework therefore targets a class of adenosines that are absent from current catalogs but resemble known m^6^A sites in sequence and regulatory context. The multi-site SELECT validation and the dCasRx-ALKBH5 perturbation are useful in this framework because they move model output from prediction toward experimentally testable regulatory hypotheses.

A recurring difficulty in m^6^A prediction is that public resources remain incomplete, heterogeneous and context dependent, since some annotations derive from broad methylation-enriched regions and others from single-nucleotide evidence^7,8,27^. Absence from a database should therefore not be read as confirmed absence of methylation. Building on this view, M6AFormer treats unannotated adenosines as candidates that require prioritization rather than as negatives by default, so the framework complements experimental mapping rather than replacing it. This choice matters because negative labels in computational m^6^A studies often reflect lack of annotation rather than experimentally confirmed absence of methylation^28,29^. To keep interpretation cautious, we separated three groups throughout the analysis, namely annotated m^6^A sites, unannotated high-confidence candidates, and a random adenosine background, which avoids equating unobserved sites

The interpretation of unannotated high-confidence candidates rests on convergent evidence at two levels. At the model level, the attribution signal centered on the candidate adenosine and its surrounding context indicates that predictions are grounded in recognizable m^6^A sequence grammar, and the performance drop after removing the transformer indicates that broader sequence context also contributes. This fits the view that m^6^A deposition reflects not a short motif alone but a combination of local sequence, transcript architecture and regulatory context^9,10,12^. At the biological level, unannotated high-confidence candidates were distributed around transcript landmarks in the expected manner, were enriched near writers, regulators and readers, and were over-represented in RNA-processing, translation, neuronal, cell-cycle and adhesion-related pathways, consistent with established roles of m^6^A in RNA metabolism and cell-state regulation^30,31^. They also overlapped RMVar and RMDisease annotations and showed enrichment in S-LDSC and GARFIELD analyses, which suggests that these candidates may mark regions enriched for disease-associated regulatory variation not fully represented in current m^6^A annotations. These analyses are not redundant with the benchmark, since they test whether prioritized candidates connect to biological programs and genetic variation. Together, they do not establish that every unannotated high-confidence candidate is methylated or functional. Rather, they show that sequence-prioritized adenosines are enriched for the molecular and genetic features expected of candidate regulatory m^6^A sites.

The NEU4 perturbation provides a focused example of how this framework can generate functional hypotheses. Targeted dCasRx-ALKBH5 editing reduced local m^6^A abundance, decreased NEU4 transcript level and altered migration and invasion phenotypes without a detectable proliferation effect. These data suggest that the prioritized m^6^A site contributes to transcript regulation and cell behavior in HepG2 cells. The result should be read as evidence for functional relevance rather than a complete mechanism. The direction of the expression change is consistent with a model in which local m^6^A contributes positively to NEU4 RNA abundance, possibly through reader-mediated stabilization or m^6^A-dependent changes in RNA-protein interactions^32,33^. The responsible reader proteins, downstream RNA-fate changes and the direct link between NEU4 expression and the migration and invasion phenotype remain to be determined. The broader point is that M6AFormer can nominate sites suitable for such mechanistic follow-up, and not that a single perturbation fully defines the regulatory pathway.

Several limitations should be considered. Computational predictions are not direct measurements of methylation, so even unannotated high-confidence candidates require experimental validation and may include false positives or condition-specific sites not methylated in every cell type. The transcriptome-wide scan and the head-to-head benchmark also relied on different model configurations. Because genome-wide scoring evaluates every adenosine, most of which lie outside the DRACH motif, we used the all-adenosine model for discovery, since its negative training distribution matches the genomic adenosine population, whereas the DRACH-restricted model was suited to the motif-controlled comparison against existing predictors. One trade-off is that the all-adenosine model separates methylated from unmethylated adenosines within the motif less sharply than the DRACH-restricted model. We limited this by using a stringent probability cutoff and by validating high-confidence candidates with orthogonal RIP-seq enrichment and site-specific assays that do not depend on the scanning model. Benchmark performance itself depends on training data, negative-site definitions and evaluation design. Each comparator was confirmed to reproduce its published performance on its own held-out data before being benchmarked here, so the margin we report reflects cross-dataset generalization rather than a faulty re-implementation, and continued community benchmarking on independent single-nucleotide maps will be valuable. The disease analyses are supportive but not causal, since RMVar and RMDisease overlap, S-LDSC and GARFIELD connect candidates to disease-associated variation at the level of association rather than direct evidence that specific m^6^A changes drive disease risk^34^. Finally, the dCasRx-ALKBH5 experiment validates one prioritized site in one cellular context, and additional sites, cell types, rescue experiments and reader perturbations will be needed to define generality and mechanism.

Looking ahead, several directions will be important for turning candidate m^6^A predictions into biological understanding. A first priority is the generation of higher-resolution and quantitative single-nucleotide m^6^A maps across many cell types and conditions, which would provide more reliable training and evaluation data than current heterogeneous resources. A second priority is to move from binary calls toward models of methylation stoichiometry, so that predictions reflect how much a site is modified rather than only whether it is modified. A related challenge is to separate constitutive m^6^A sites from context-dependent regulatory events, which will require modeling cell-state and condition-specific regulation rather than a single static landscape. Beyond sequence, integrating RNA secondary structure together with reader and writer binding may help explain why particular adenosines are selected for methylation. Closing the gap between association and causation will also be essential, since large-scale functional and perturbation studies are still needed to establish which predicted sites carry regulatory or disease relevance. Progress on these fronts would allow sequence-based prioritization to contribute more directly to a mechanistic and clinically interpretable view of the m^6^A epitranscriptome.

## Material and methods

### Dataset preparation and processing

Base-resolution m^6^A sites were compiled from m6A-Atlas (427,760 sites) and RMBase (879,436), unified to hg38 and merged into a non-redundant reference of 1,044,601 sites. To keep benchmarking blind to competing tools, sequences overlapping the training data of the three benchmarked predictors including MST-m6A and CLSM6A (123,822 shared positives) and deepSRAMP (246,103 sites) were removed before partitioning. After sequence-level deduplication, the 694,578 remaining positive sites were split 95:5 into training and an independent held-out test set; per-configuration counts are given in Supplementary Table S10.

Negatives were non-m^6^A adenosines of two types: DRACH-matched (motif-conforming) and random, drawn at same number of the positive count. Two window lengths, 201 nt and 801 nt (±100 and ±400 nt around the central adenosine), were compared; windows truncated at transcript boundaries were N-padded.

### Model architecture and training

All models were implemented in PyTorch v2.12.1 and trained by minimizing a binary cross-entropy loss (BCEWithLogitsLoss) with the AdamW optimizer (initial learning rate 1 × 10^-4^, weight decay 1 × 10^-3^) and a batch size of 64. The learning rate followed a warm-up cosine schedule (warm-up fraction 0.1, minimum learning-rate ratio 0.01). For each configuration, the training partition was further divided into fitting and inner-validation subsets at a 90:10 ratio using a stratified, sequence-grouped split; for the main all-adenosine 801-nt model (ALL-801), which achieved the best overall performance and was used for the subsequent transcriptome-wide scan, this yielded 1,187,730 fitting sites and 131,970 inner-validation sites. Architectural hyper-parameters (latent dimension 128, four attention heads, two transformer layers, dropout 0.1) were fixed after selection on the internal validation set (Supplementary Table S11).

### Benchmarking against existing m^6^A predictors

M6AFormer was compared with deepSRAMP^24^, CLSM6A^18^, MST-m6A^23^ and a CNN baseline on the complete independent held-out test set, comprising the held-out positive m^6^A sites and an equal number of DRACH-motif-restricted non methylated sites as negatives. Each tool received input in its native format, and sequences overlapping training data were removed before scoring. Before benchmarking, each comparator was verified to reproduce its published performance on its own held-out data (deepSRAMP, AUROC 0.959 versus 0.967 reported; MST-m6A, accuracy 0.6098 to 0.8855, matching its reported range; Supplementary Table S5), so that any difference observed on our benchmark reflects generalization rather than an incorrect re-implementation. Each comparator was represented by its best-performing released model on this site set, namely T9 for CLSM6A and T5 for MST-m6A. Comparators were selected on the basis of three criteria, namely availability of source code, a pre-trained model, and clearly defined training and test datasets (Supplementary Table S1). Performance was quantified with scikit-learn v1.9.0 using standard classification metrics, including AUROC, AUPRC, accuracy, precision, sensitivity, specificity, and MCC.

To determine whether the higher discrimination of M6AFormer was statistically significant rather than attributable to sampling variation, we compared ROC curves on this common held-out set using the DeLong test for two correlated areas under the curve, as implemented in pROC v1.18.5^35^ (roc.test, method = “delong”). Because all methods were scored on the identical set of sites, we used the paired formulation for correlated ROC curves. M6AFormer, in the DRACH-restricted 801-nt configuration, served as the reference model and was contrasted pairwise against CLSM6A, MST-m6A and deepSRAMP, with each comparator represented by its best-performing configuration. All pairwise differences were significant by two-sided DeLong test.

To evaluate performance under matched training and testing conditions, each model was assessed with its own cell-line-specific model on the corresponding cell-line subset of the held-out data, alongside M6AFormer in the DRACH-restricted 801-nt configuration and the CNN baseline. As in the pooled benchmark, negatives were DRACH-motif-restricted non-methylated adenosines drawn from the same transcripts. Discrimination was summarized by AUROC with 95% confidence intervals estimated from 1,000 bootstrap resamples, together with the MCC. Cell lines for which a method provided no dedicated model or lacked test data were omitted, so that only cell lines with an available model and test data were compared.

### Ablation and interpretation

Ablation analysis was performed to evaluate the contribution of major architectural components. The full model was compared with variants lacking the transformer encoder, multi-scale convolution module, residual convolution blocks or attention-pooling component. Ablated models used the same input windows, training data, validation strategy and evaluation metrics as the full model. Performance differences were summarized by AUROC, AUPRC, accuracy and Matthews correlation coefficient on held-out evaluation sets.

Feature attribution analysis was used to determine whether model predictions were supported by biologically plausible sequence features. Attribution scores were calculated with Integrated Gradients in Captum v0.9.0^36^ used for the feature-importance panels. A zero-valued one-hot tensor was used as the baseline, and attributions were computed with 50 integration steps. Position-wise importance was defined as the mean absolute attribution summed across nucleotide channels.

### Transcriptome-wide candidate-site scanning

Transcriptome-wide scanning was performed with the all-adenosine 801-nt model (ALL-801). Genome-wide discovery scores every candidate adenosine rather than only motif-restricted sites, so the model trained with all-adenosine negatives matched this setting and was adopted as the primary M6AFormer model for discovery. Within the all-adenosine setting, the 801-nt window gave stronger held-out performance than the 201-nt window. For transcriptome-wide discovery, all eligible adenosine-centered sites in annotated exonic transcript regions were enumerated from the hg38 reference genome. On the positive strand, genomic A bases were considered candidate transcriptomic adenosines. On the negative strand, genomic T bases were treated as transcriptomic adenosines after strand correction. All retained candidate sites were scored with the ALL-801 checkpoint in PyTorch. Predicted sites were annotated by chromosome, position, strand, transcript region, gene symbol, DRACH status and overlap with m6A-Atlas or RMBase. Sites overlapping the curated m^6^A database union were labeled annotated, whereas sites lacking overlap with this union were labeled unannotated.

Candidate sites were summarized at multiple probability thresholds. To distinguish high-confidence predictions from lower-confidence calls, the probability cutoff was chosen by jointly examining, on the held-out test set, precision and sensitivity as a function of the threshold, the agreement between predicted probabilities and observed positive rates (calibration), and the number of candidate sites retained as the threshold was tightened. A cutoff of prob ≥ 0.99 was adopted as the primary high-confidence operating point, selected to maximize precision while preserving calibration and retaining a substantial candidate set. This threshold was used to define the high-confidence unannotated predictions analysed throughout. Gene-level summaries counted genes containing at least one candidate site in each category. Metagene profiles were generated by assigning sites to 5’ UTR, coding sequence and 3’ UTR bins and comparing the distribution of annotated sites with high-confidence unannotated predictions.

### Regulatory-context analysis of m^6^A machinery

Public RNA immunoprecipitation profiling (RIP-seq) datasets were used to evaluate whether high-confidence M6AFormer candidates were enriched in regulatory contexts associated with m^6^A machinery and m^6^A-related RNA-binding proteins. Targets included IGF2BP1^33^, IGF2BP2^33^, IGF2BP3^33^, METTL3^37^, METTL14^37^, METTL16^37^, RBM15^38^ and VIRMA^39^.

Reads were adapter- and quality-trimmed with Trim Galore v0.6.11^40^ using quality cutoff 15, minimum length 15 and stringency 3, with FastQC v0.12.1^41^ reports generated after trimming. Reads were aligned to the hg38 index with STAR v2.7.11b^42^ in two-pass mode, and sorted BAM files were indexed with samtools v1.23.1^43^. Processed binding or enrichment intervals for each target were intersected with annotated and unannotated M6AFormer candidate sets. Overlap rates and local enrichment patterns were then compared and visualized between annotated sites, high-confidence unannotated sites and background candidate sites using pyGenomeTrack v3.9^44^.

### Functional enrichment and disease-associated variant analyses

Functional enrichment of genes harboring candidate m^6^A sites was performed with clusterProfiler v4.14.6^45^ and DOSE v4.0.0^46^, testing KEGG pathways, GO Biological Process terms, Disease Ontology (DO) and DisGeNet (DGN) against a background of all mappable genes from the scanned transcriptome. P values were adjusted by the Benjamini-Hochberg method and terms with q < 0.05 were considered significant. Overlap with m^6^A-associated disease variants was tested using RMVar^47^ and RMDisease^48^. Variant records were converted to strand-aware site identifiers, and exact and local-window overlaps for annotated sites, unannotated sites and matched background sets were compared with Fisher’s exact tests. Trait-level enrichment was evaluated with S-LDSC v1.0.1^49^ and GARFIELD v1.34.0^50^. Candidate-site were expanded by 500 bp on each side and converted to annotations against 1000 Genomes European reference SNPs, and analyses were run over ten GWAS traits (Supplementary Table S12)^49,51,52^.

### Cell culture

HepG2 cells were maintained in Dulbecco’s modified Eagle’s medium (DMEM) supplemented with 1% penicillin-streptomycin and 10% fetal bovine serum at 37 °C in a humidified incubator with 5% CO_2_.

### SELECT

Site-specific m^6^A abundance was measured with SELECT as previously described^53^. Total RNA was extracted from control and dCasRx-ALKBH5-treated cells using RNeasy Mini Kit (QIAGEN, 74106). SELECT reactions used site-specific upstream, downstream and qPCR primers designed around each candidate adenosine. The assay included multiple model-prioritized sites and positive and negative validation categories. Relative m^6^A abundance was calculated from qPCR readouts and normalized to the corresponding control condition. Sequences should be provided in Supplementary Table S13.

### m^6^A site-specific perturbation

The pMSCV-dCasRx-ALKBH5-PURO construct was a gift from Qi Xie (Addgene plasmid # 175582; http://n2t.net/addgene:175582; RRID:Addgene_175582)^54^. Target gRNAs were designed using the CRISPOR^55^, and the scramble gRNA was as described by Xia et al.^54^ (Supplementary Table S13). Each guide RNA, together with the upstream U6 promoter and downstream stem sequence, was synthesized by IDT and ligated into the pCR-Blunt II-TOPO vector (Invitrogen, K280020). Single colonies were confirmed by Sanger sequencing for correct orientation and absence of mutation. HepG2 cells were electroporated with a total of 3 µg of dCasRx-ALKBH5 and gRNA vector using the Cell Line Nucleofector Kit V (Lonza, VCA-1003) per the manufacturer’s protocol, and transfection efficiency was assessed at 12 h

### RT-qPCR, CCK8 and transwell assays

NEU4 mRNA expression after perturbation was measured by RT-qPCR. Total RNA was reverse-transcribed into cDNA using the LunaScript RT SuperMix Kit (NEB, E3010). Gene expression was quantified by qRT-PCR using PowerTrack SYBR Green Master Mix (Applied Biosystems, A46110), and relative expression levels were normalized to GAPDH (Supplementary Table S13). Each group comprised three biological replicates and groups were compared by two-sided Student’s t-test.

Cell proliferation was measured by Cell Counting Kit - 8 assay (Sigma-Aldrich, 96992), with corrected OD450 recorded at 24, 48 and 96 h and compared between control and dCasRx-ALKBH5-treated groups at each time point.

Cell migration and invasion were assessed by transwell assays. Briefly, cells were seeded into 8-µm transwell inserts, uncoated for migration (Corning, 353097) and Matrigel-coated for invasion (Corning, 354480). After incubation for 16 h (migration) and 36 h (invasion), non-migrated cells on the upper surface were removed with a cotton swab, and the inserts were washed twice with PBS, fixed in 4% paraformaldehyde for 30 min and stained with crystal violet for 20 min. Migrated and invaded cells were imaged on an EVOS cell-imaging system (Invitrogen) and counted separately for the two assays using ImageJ^56^.

### Statistical analysis and reproducibility

For model evaluation, n denotes the number of candidate sites in the corresponding test set. For wet-lab experiments, n denotes independent biological replicates unless otherwise specified. AUROC confidence intervals were estimated by bootstrap resampling. Functional enrichment analyses used Benjamini-Hochberg-adjusted q-value < 0.05 as the significance threshold. Disease-variant overlap analyses used Fisher’s exact tests.

## Acknowledgements

This work was supported by Deutsche Forschungsgemeinschaft (DFG, German Research Foundation), EXC 2026, Cardio-Pulmonary Institute (CPI), Project ID 390649896.

## Author contributions

**Zhixin Niu**: Conceptualization, Data curation, Formal analysis, Investigation, Methodology, Software, Validation, Visualization, Writing - original draft, Writing - review & editing. **Chang Liu**: Investigation, Validation, Writing - review & editing. **Lei Gu**: Conceptualization, Funding acquisition, Project administration, Resources, Supervision, Writing - review & editing.

## Data availability

The high-confidence transcriptome-wide m^6^A prediction table and the GO, KEGG, DO and DGN enrichment tables generated in this study have been deposited at Zenodo under https://doi.org/10.5281/zenodo.2129630657. Model checkpoints, candidate-site BED files and processed prediction tables are available in the same repository. The M6AFormer source code is available at GitHub (https://github.com/zhixinniu/M6AFormer)^58^ and is released under the MIT license. The trained model is additionally distributed as a package via PyPI (https://pypi.org/project/m6aformer/0.1.2/). This study analysed the following publicly available resources. m^6^A site annotations were obtained from m6A-Atlas v2.0^7^ and RMBase v3.0^8^. m^6^A-associated disease-variant annotations were obtained from RMVar 2.0^47^ and RMDisease^48^. GWAS summary statistics used for the stratified LD-score regression and GARFIELD analyses, with their accession identifiers and sources, are listed in Supplementary Table S12. NEU4 expression across cell lines was obtained from the Human Protein Atlas^59^, and NEU4 tumour-versus-normal expression from the TCGA-LIHC cohort^60^. Public RIP-seq datasets used to profile writer, regulator and reader binding around predicted m^6^A sites were obtained from the Gene Expression Omnibus under accessions GSE90639 (IGF2BP1/2/3)^33^, GSE156797 (METTL3, METTL14 and METTL16)^37^, GSE73893 (RBM15 RIP-seq)^38^, GSE102493 (VIRMA)^39^. Datasets used for model training and benchmarking, and the comparator datasets for CLSM6A^18^, MST-m6A^23^ and deepSRAMP^24^, are derived from the public resources above.

## References

1 Baquero-Pérez, B. et al. N6-methyladenosine modification is not a general trait of viral RNA genomes. Nature Communications 15, 1964, doi:10.1038/s41467-024-46278-9 (2024).

2 Dierks, D. et al. Passive shaping of intra- and intercellular m6A dynamics via mRNA metabolism. eLife 13, RP100448, doi:10.7554/eLife.100448 (2025).

3 Vaid, R. et al. METTL3 drives telomere targeting of TERRA lncRNA through m6A-dependent R-loop formation: a therapeutic target for ALT-positive neuroblastoma. Nucleic Acids Research 52, 2648–2671, doi:10.1093/nar/gkad1242 (2024).

4 Lee, E. S., et al. *N*-6-methyladenosine (m6A) promotes the nuclear retention of mRNAs with intact 5’ splice site motifs. Life Science Alliance 8, e202403142, doi:10.26508/lsa.202403142 (2025).

5 Zhou, Y. et al. m6A sites in the coding region trigger translation-dependent mRNA decay. Molecular Cell 84, 4576–4593.e4512, 10.1016/j.molcel.2024.10.033 (2024).

6 Vandelli, A., Broglia, L., Armaos, A., Delli Ponti, R. & Tartaglia, G. G. Rationalizing the effects of RNA modifications on protein interactions. Molecular Therapy Nucleic Acids 35, 102391, 10.1016/j.omtn.2024.102391 (2024).

7 Liang, Z. et al. m6A-Atlas v2.0: updated resources for unraveling the N6-methyladenosine (m6A) epitranscriptome among multiple species. Nucleic Acids Research 52, D194–D202, doi:10.1093/nar/gkad691 (2024).

8 Xuan, J. et al. RMBase v3.0: decode the landscape, mechanisms and functions of RNA modifications. Nucleic Acids Research 52, D273–D284, doi:10.1093/nar/gkad1070 (2024).

9 Wang, Y., Wang, S., Meng, Z., Liu, X.-M. & Mao, Y. Determinant of m6A regional preference by transcriptional dynamics. Nucleic Acids Research 52, 3510–3521, doi:10.1093/nar/gkae169 (2024).

10 Shachar, R. et al. Dissecting the sequence and structural determinants guiding m6A deposition and evolution via inter- and intra-species hybrids. Genome Biology 25, 48, doi:10.1186/s13059-024-03182-1 (2024).

11 Santos-Rodriguez, G. et al. Isoform-specific m6A deposition and coordinated splicing shape mammalian transcriptome evolution. Nature Communications 17, 5075, doi:10.1038/s41467-026-72124-1 (2026).

12 Uzonyi, A. et al. Exclusion of m6A from splice-site proximal regions by the exon junction complex dictates m6A topologies and mRNA stability. Molecular Cell 83, 237–251.e237, 10.1016/j.molcel.2022.12.026 (2023).

13 Tang, Y. et al. m6A-Atlas: a comprehensive knowledgebase for unraveling the N6-methyladenosine (m6A) epitranscriptome. Nucleic Acids Research 49, D134–D143, doi:10.1093/nar/gkaa692 (2021).

14 Meyer, Kate D. et al. Comprehensive Analysis of mRNA Methylation Reveals Enrichment in 3’ UTRs and near Stop Codons. Cell 149, 1635–1646, 10.1016/j.cell.2012.05.003 (2012).

15 McIntyre, A. B. R. et al. Limits in the detection of m6A changes using MeRIP/m6A-seq. Scientific Reports 10, 6590, doi:10.1038/s41598-020-63355-3 (2020).

16 Dominissini, D., Moshitch-Moshkovitz, S., Amariglio, N. & Rechavi, G. in Methods in Enzymology Vol. 560 (ed Chuan He) 131–147 (Academic Press, 2015).

17 Shen, W. et al. GLORI for absolute quantification of transcriptome-wide m6A at single-base resolution. Nature Protocols 19, 1252–1287, doi:10.1038/s41596-023-00937-1 (2024).

18 Zhang, Y. et al. Interpretable prediction models for widespread m6A RNA modification across cell lines and tissues. Bioinformatics 39, doi:10.1093/bioinformatics/btad709 (2023).

19 Pratanwanich, P. N. et al. Identification of differential RNA modifications from nanopore direct RNA sequencing with xPore. Nature Biotechnology 39, 1394–1402, doi:10.1038/s41587-021-00949-w (2021).

20 Hendra, C. et al. Detection of m6A from direct RNA sequencing using a multiple instance learning framework. Nature Methods 19, 1590–1598, doi:10.1038/s41592-022-01666-1 (2022).

21 Zhong, Z.-D. et al. Systematic comparison of tools used for m6A mapping from nanopore direct RNA sequencing. Nature Communications 14, 1906, doi:10.1038/s41467-023-37596-5 (2023).

22 Zhou, Y., Zeng, P., Li, Y.-H., Zhang, Z. & Cui, Q. SRAMP: prediction of mammalian N6-methyladenosine (m6A) sites based on sequence-derived features. Nucleic Acids Research 44, e91–e91, doi:10.1093/nar/gkw104 (2016).

23 Su, Q., Phan, L. T., Pham, N. T., Wei, L. & Manavalan, B. MST-m6A: A Novel Multi-Scale Transformer-based Framework for Accurate Prediction of m6A Modification Sites Across Diverse Cellular Contexts. Journal of Molecular Biology 437, 168856, 10.1016/j.jmb.2024.168856 (2025).

24 Fan, R. et al. A combined deep learning framework for mammalian m6A site prediction. Cell Genomics 4, 100697, 10.1016/j.xgen.2024.100697 (2024).

25 Song, R., J Sutton, G., Li, F., Liu, Q. & Wong, J. J.-L. Variable calling of m6A and associated features in databases: a guide for end-users. Briefings in Bioinformatics 25, doi:10.1093/bib/bbae434 (2024).

26 Zhang, X. et al. NEU4 inhibits motility of HCC cells by cleaving sialic acids on CD44. Oncogene 40, 5427–5440, doi:10.1038/s41388-021-01955-7 (2021).

27 Zhao, X. et al. m6AConquer: a consistently quantified and orthogonally validated database for the N6-methyladenosine (m6A) epitranscriptome. Nucleic Acids Research 54, D204–D218, doi:10.1093/nar/gkaf1204 (2026).

28 Yang, J. et al. Machine learning–augmented m6A-Seq analysis without a reference genome. Briefings in Bioinformatics 26, doi:10.1093/bib/bbaf235 (2025).

29 Wang, L. et al. Block sparse Bayes-based fuzzy system for RNA N6-methyladenosine sites prediction. PLOS Computational Biology 21, e1013621, doi:10.1371/journal.pcbi.1013621 (2025).

30 Wei, J. & He, C. RNA modifications in gene regulation: Functions and pathways. Cell 189, 1591–1619, doi:10.1016/j.cell.2026.01.006 (2026).

31 Zhou, Y., Cao, P. & Zhu, Q. The regulatory role of m6A in cancer metastasis. Frontiers in Cell and Developmental Biology Volume 13 - 2025, doi:10.3389/fcell.2025.1539678 (2025).

32 Gao, B. et al. RNA demethylase ALKBH5 regulates cell cycle progression in DNA damage response. Scientific Reports 15, 16059, doi:10.1038/s41598-025-01207-8 (2025).

33 Huang, H. et al. Recognition of RNA N6-methyladenosine by IGF2BP proteins enhances mRNA stability and translation. Nature Cell Biology 20, 285–295, doi:10.1038/s41556-018-0045-z (2018).

34 Liu, S., Zhao, X., Wei, Z., Carr, D. F. & Moraros, J. Decoding causal m6A: a bioinformatics roadmap for psychiatric disorders. Briefings in Bioinformatics 27, doi:10.1093/bib/bbag251 (2026).

35 Robin, X. et al. pROC: an open-source package for R and S+ to analyze and compare ROC curves. BMC Bioinformatics 12, 77, doi:10.1186/1471-2105-12-77 (2011).

36 Kokhlikyan, N., et al. Captum: A unified and generic model interpretability library for pytorch. *arXiv preprint arXiv:2009.07896* (2020).

37 Su, R. et al. METTL16 exerts an m6A-independent function to facilitate translation and tumorigenesis. Nature Cell Biology 24, 205–216, doi:10.1038/s41556-021-00835-2 (2022).

38 Zhang, L. et al. Cross-talk between PRMT1-mediated methylation and ubiquitylation on RBM15 controls RNA splicing. eLife 4, e07938, doi:10.7554/eLife.07938 (2015).

39 Yue, Y. et al. VIRMA mediates preferential m6A mRNA methylation in 3’ UTR and near stop codon and associates with alternative polyadenylation. Cell Discovery 4, 10, doi:10.1038/s41421-018-0019-0 (2018).

40 Krueger, F. J., Frankie; Ewels, Phil; Afyounian, Ebrahim; Schuster-Boeckler, Benjamin. FelixKrueger/TrimGalore, <https://zenodo.org/records/5127899> (2021).

41 Andrews, S. FastQC: A Quality Control Tool for High Throughput Sequence Data, <https://www.bioinformatics.babraham.ac.uk/projects/fastqc/> (2010).

42 Dobin, A. et al. STAR: ultrafast universal RNA-seq aligner. Bioinformatics 29, 15–21, doi:10.1093/bioinformatics/bts635 (2013).

43 Danecek, P. et al. Twelve years of SAMtools and BCFtools. GigaScience 10, doi:10.1093/gigascience/giab008 (2021).

44 Lopez-Delisle, L., et al. pyGenomeTracks: reproducible plots for multivariate genomic datasets Bioinformatics 37, 422–423, doi:10.1093/bioinformatics/btaa692 (2021).

45 Yu, G., Wang, L. G., Han, Y. & He, Q. Y. clusterProfiler: an R package for comparing biological themes among gene clusters. Omics 16, 284–287, doi:10.1089/omi.2011.0118 (2012).

46 Yu, G., Wang, L.-G., Yan, G.-R. & He, Q.-Y. DOSE: an R/Bioconductor package for disease ontology semantic and enrichment analysis. Bioinformatics 31, 608–609, doi:10.1093/bioinformatics/btu684 (2015).

47 Huang, Y. et al. RMVar 2.0: an updated database of functional variants in RNA modifications. Nucleic Acids Research 53, D275–D283, doi:10.1093/nar/gkae924 (2025).

48 Chen, K. et al. RMDisease: a database of genetic variants that affect RNA modifications, with implications for epitranscriptome pathogenesis. Nucleic Acids Research 49, D1396–D1404, doi:10.1093/nar/gkaa790 (2021).

49 Finucane, H. K. et al. Partitioning heritability by functional annotation using genome-wide association summary statistics. Nature Genetics 47, 1228–1235, doi:10.1038/ng.3404 (2015).

50 Iotchkova, V. et al. GARFIELD classifies disease-relevant genomic features through integration of functional annotations with association signals. Nature Genetics 51, 343–353, doi:10.1038/s41588-018-0322-6 (2019).

51 Bulik-Sullivan, B. K. et al. LD Score regression distinguishes confounding from polygenicity in genome-wide association studies. Nature Genetics 47, 291–295, doi:10.1038/ng.3211 (2015).

52 Gazal, S. et al. Linkage disequilibrium–dependent architecture of human complex traits shows action of negative selection. Nature Genetics 49, 1421–1427, doi:10.1038/ng.3954 (2017).

53 Xiao, Y. et al. An Elongation- and Ligation-Based qPCR Amplification Method for the Radiolabeling-Free Detection of Locus-Specific N6-Methyladenosine Modification. Angewandte Chemie International Edition 57, 15995–16000, 10.1002/anie.201807942 (2018).

54 Xia, Z. et al. Epitranscriptomic editing of the RNA N6-methyladenosine modification by dCasRx conjugated methyltransferase and demethylase. Nucleic Acids Research 49, 7361–7374, doi:10.1093/nar/gkab517 (2021).

55 Concordet, J.-P. & Haeussler, M. CRISPOR: intuitive guide selection for CRISPR/Cas9 genome editing experiments and screens. Nucleic Acids Research 46, W242–W245, doi:10.1093/nar/gky354 (2018).

56 Schneider, C. A., Rasband, W. S. & Eliceiri, K. W. NIH Image to ImageJ: 25 years of image analysis. Nature Methods 9, 671–675, doi:10.1038/nmeth.2089 (2012).

57 Niu, Z. Dataset for “M6AFormer Prioritizes Unannotated Functional m6A Candidate Sites in the Human m6A Epitranscriptome”, <10.5281/zenodo.21296306> (2026).

58 Niu, Z. M6AFormer: deep learning toolkit for transcriptome-wide N6-methyladenosine (m6A) site prediction, built on a hybrid CNN + lightweight Transformer architecture. GitHub (2026).

59 Uhlén, M. et al. Tissue-based map of the human proteome. Science 347, 1260419, doi:10.1126/science.1260419 (2015).

60 McLendon, R. et al. Comprehensive genomic characterization defines human glioblastoma genes and core pathways. Nature 455, 1061–1068, doi:10.1038/nature07385 (2008).

