## Supplementary Information for "M6AFormer Prioritizes Unannotated Functional m^6^A Candidate Sites in the Human Epitranscriptome"

**Table of contents**

### I. Supplementary Figures

Figure S1. Cell-line matched benchmarking of M6AFormer against existing predictors

Figure S2. Overlap of the full M6AFormer predicted set with RMBase and m6A-Atlas

Figure S3. Selection of the high-confidence probability threshold

Figure S4. Disease Ontology and DisGeNET enrichment of annotated and unannotated predicted sites

Figure S5. NEU4 expression across human cell lines

Figure S6. NEU4 downregulation in liver hepatocellular carcinoma

### II. Supplementary Tables

Table S1. Survey of published m6A prediction tools

Table S2. Performance of M6AFormer across four training configurations

Table S3. Performance of existing $m^6A$ predictors and the CNN baseline

Table S4. DeLong tests comparing M6AFormer with the comparator models

Table S6. Cell line–matched evaluation of M6AFormer and comparator models in four cell lines

Table S7. Ablation study of M6AFormer architectural components

Table S8. S-LDSC heritability enrichment of predicted $m^6A$ sites across traits

Table S9. GARFIELD enrichment of predicted $m^6A$ sites across GWAS traits

Table S11. M6AFormer model architecture and training hyperparameters

Table S12. GWAS summary statistics used for S-LDSC and GARFIELD analyses

Table S13. Synthetic nucleotide sequences used in this study

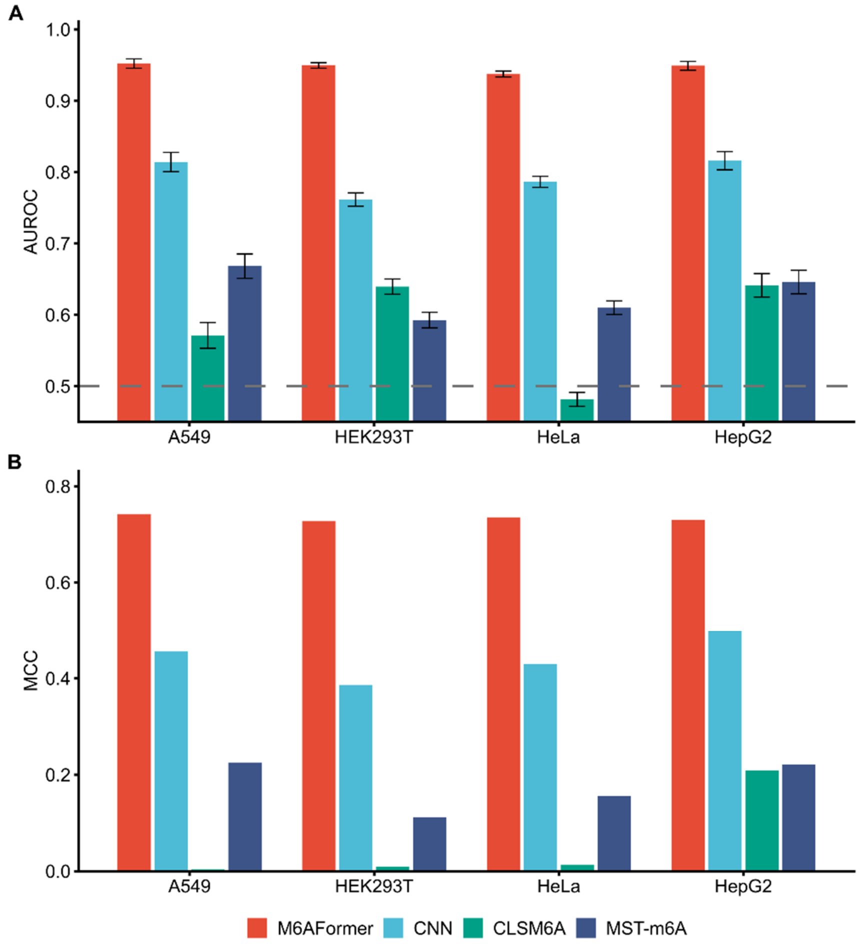


**Figure S1. Cell-line matched benchmarking of M6AFormer against existing predictors.** Each method was evaluated with its own cell-line-specific model on the corresponding cell-line subset of the held-out data, alongside M6AFormer and the CNN baseline. **(A)** AUROC for M6AFormer, CLSM6A, MST-m6A, and the CNN baseline across A549, HEK293T, HeLa and HepG2. Error bars show 95% confidence intervals from bootstrap resampling and the dashed line marks AUROC = 0.5. **(B)** MCC for the same models and cell lines. Cell lines for which a method provided no dedicated model or lacked test data are omitted.

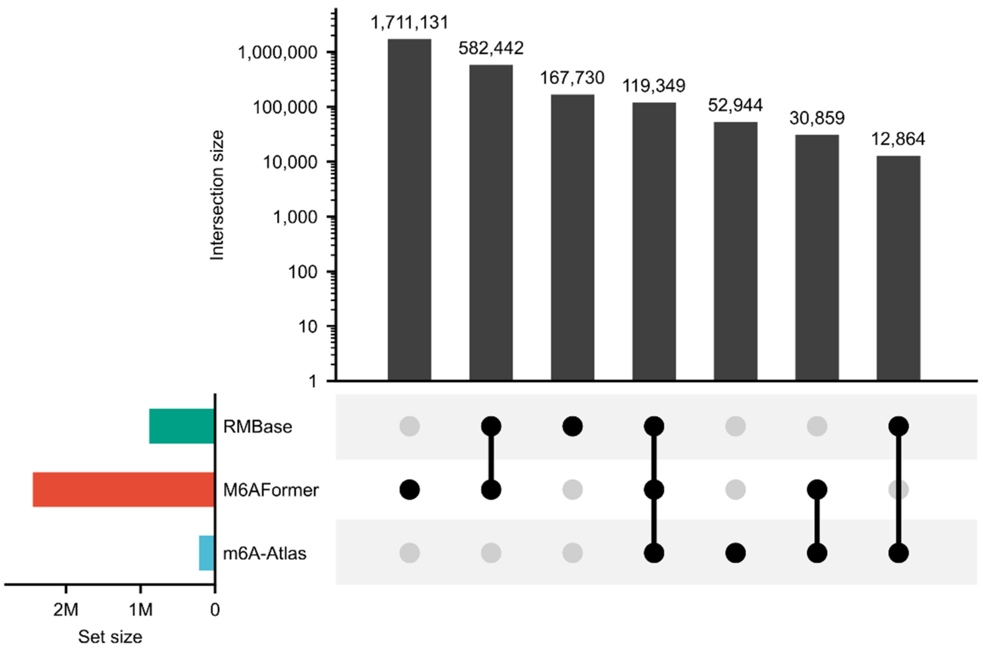


**Figure S2. Overlap of the full M6AFormer predicted set with existing m$^6$A databases.** UpSet plot comparing all M6AFormer-predicted sites with RMBase and m6A-Atlas. Horizontal bars (lower left) show the size of each set. Vertical bars show intersection sizes on a log scale, with filled dots below indicating the set combination. Most predicted adenosines lie outside both databases, whereas database-supported predictions are largely shared between RMBase and m6A-Atlas.

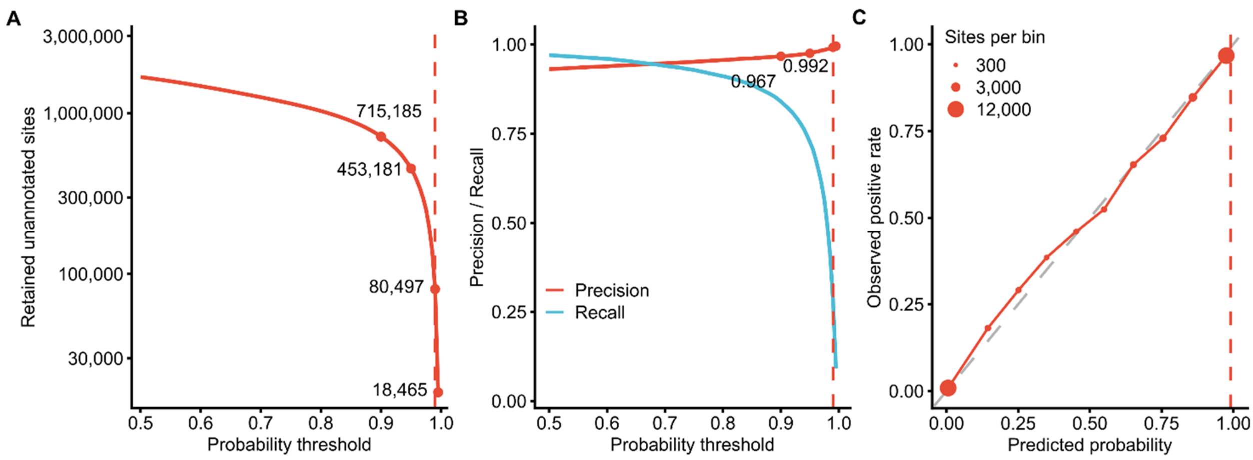


**Figure S3. Selection of the high-confidence probability threshold. (A)** Number of retained unannotated predicted sites as a function of the probability threshold (log scale), marked at 0.9, 0.95, 0.99 and 0.995. **(B)** Precision and recall on the independent held-out test set as a function of the threshold. **(C)** Calibration of predicted probabilities, showing the observed positive rate against the predicted probability on the held-out test set; the dashed diagonal denotes perfect calibration. Dotted vertical lines mark 0.9 and 0.95, and the dashed red line marks the adopted cutoff of prob > 0.99.

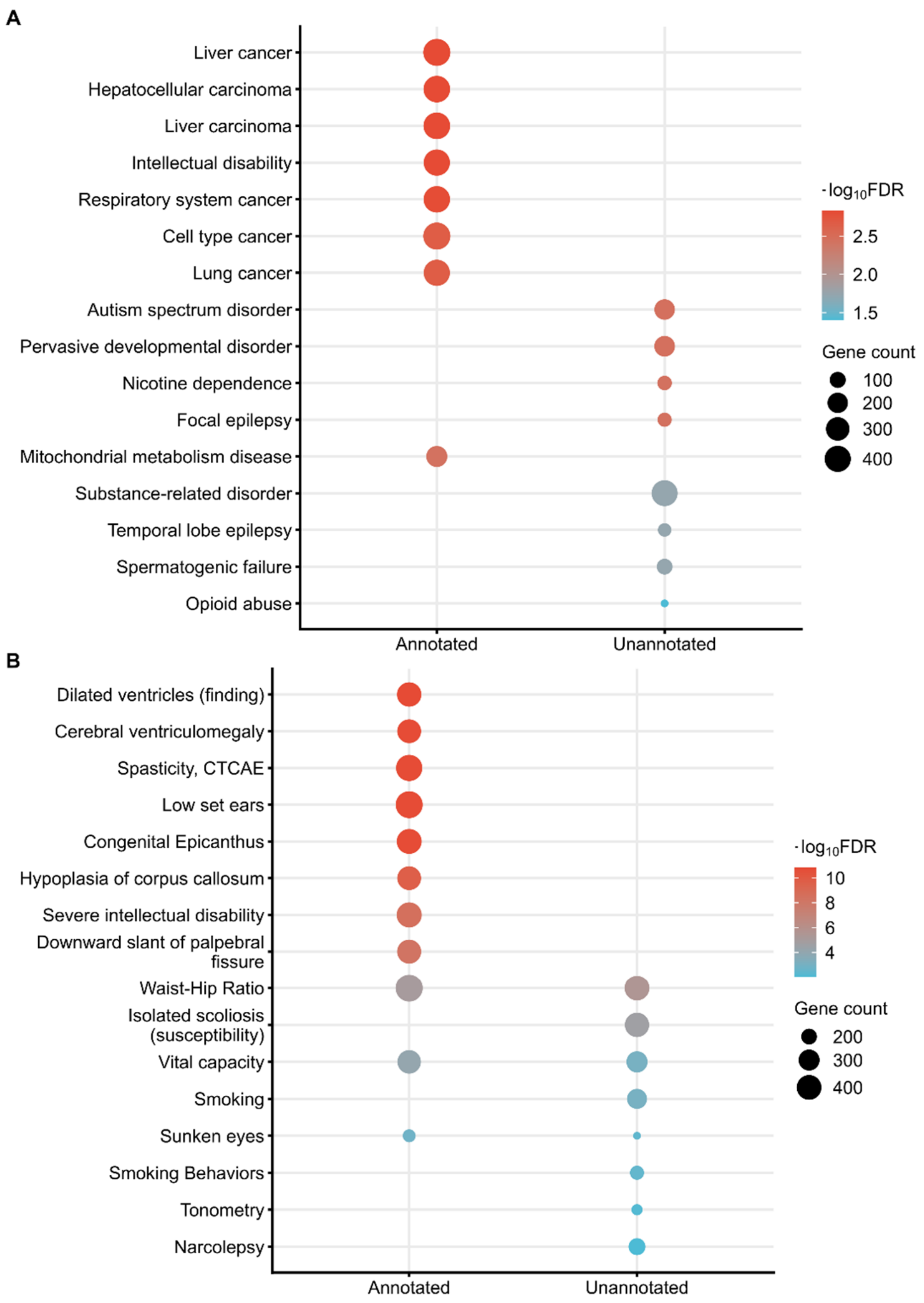


**Figure S4. Disease Ontology and DisGeNET enrichment of annotated and unannotated predicted sites.** Dot plots of the top enriched terms for genes associated with annotated and unannotated high-confidence M6AFormer predictions. **(A)** DO terms. **(B)** DGN terms. Dot size indicates the number of genes in each term and colour indicates enrichment significance (-log10 FDR).

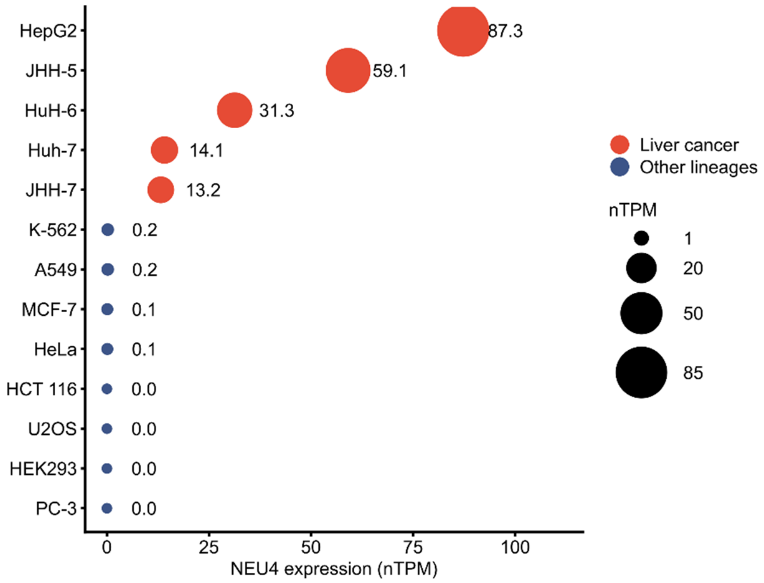


**Figure S5. NEU4 expression across human cell lines.** NEU4 mRNA abundance (nTPM) across cell lines, ordered by expression. Liver cancer lines are shown in red and other lineages in blue, with dot size proportional to nTPM. NEU4 is most highly expressed in HepG2 (87.3 nTPM), supporting its selection for site-specific perturbation in a cell line with sufficient expression for reliable detection.

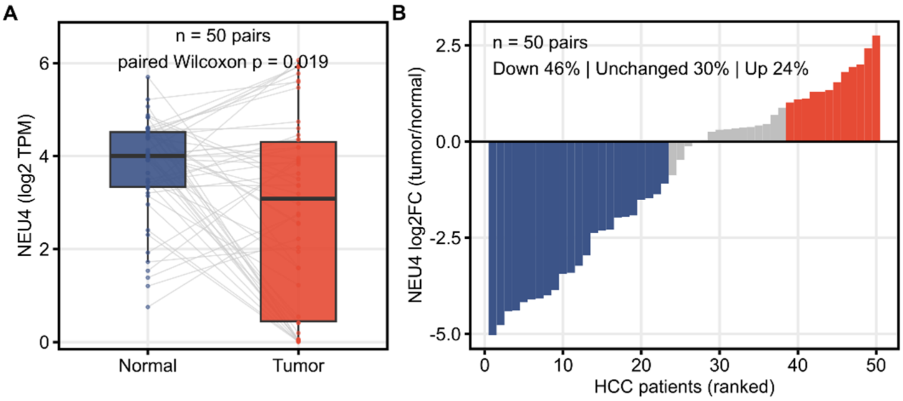


**Figure S6: NEU4 downregulation in liver hepatocellular carcinoma**. **(A)**. NEU4 expression (log2 TPM) in 50 matched pairs of liver tumour and adjacent normal liver from TCGA-LIHC. Grey lines connect matched pairs. The p value is from a two-sided paired Wilcoxon signed-rank test. **(B)** Per-pair NEU4 log2 fold change (tumour over normal) ranked across the 50 pairs. Bars are coloured as down (log2FC at most -1), unchanged, or up (log2FC at least 1).

**Table S1. Survey of published m$^6$A prediction tools**

| Tool | Architecture | Code availability | Pre-trained model | Defined train/test partition |
|---|---|---|---|---|
| m6A-SPP | CNN + Transformer | Yes | No | Yes |
| LOCO-m6A | CNN | Yes | No | Yes |
| **MST-m6A** | **Transformer** | **Yes** | **Yes** | **Yes** |
| m6A-BERT-Deg | Transformer | Yes | No | Yes |
| **deepSRAMP** | **CNN + Transformer** | **Yes** | **Yes** | **Yes** |
| **CLSM6A** | **CNN** | **Yes** | **Yes** | **Yes** |
| DL-m6A | CNN | Yes | No | Yes |
| GR-m6A | CNN | Yes | No | Yes |
| TS-m6A-DL | CNN | No | No | No |
| Rm-LR | Transformer | No | No | No |
| m6A-BERT-Stacking | RNN + Transformer | Yes | No | Yes |
| im6A-TS-CNN | CNN | Yes | Yes | No |
| m6A-NeuralTool | CNN | No | No | No |
| DeepM6ASeq | CNN | Yes | No | Yes |
| iN6-Methyl | CNN | No | No | No |
| m6A-neural-network | CNN | Yes | No | No |
| ELMo4m6A | CNN | Yes | No | Yes |
| MultiRM | CNN | Yes | Yes | No |
| iM6A | CNN | Yes | Yes | No |
| DeepPromise | CNN | No | No | No |
| BERMP | RNN | No | No | Yes |
| BLAM6A-Merge | CNN | Yes | No | Yes |
| Deepm6A-MT | CNN | No | No | No |

**Table S2. Performance of M6AFormer across four training configurations**

| Model | AUROC | AUPRC | Acc | Precision | Sen | Spe | MCC |
|---|---|---|---|---|---|---|---|
| DRACH801 | 0.9166 | 0.9161 | 0.8308 | 0.7961 | 0.8895 | 0.7722 | 0.6663 |
| DRACH201 | 0.8555 | 0.8608 | 0.7558 | 0.7092 | 0.8673 | 0.6444 | 0.5249 |
| ALL801 | 0.9839 | 0.9797 | 0.9486 | 0.9291 | 0.9714 | 0.9259 | 0.8982 |
| ALL201 | 0.9783 | 0.9720 | 0.9396 | 0.9132 | 0.9716 | 0.9077 | 0.8811 |

**Table S3. Performance of existing $m^6A$ predictors and the CNN baseline**

| Model | AUROC | AUPRC | Acc | Precision | Sen | Spe | MCC |
|---|---|---|---|---|---|---|---|
| M6AFormer (DRACH801) | 0.916583 | 0.916009 | 0.834425 | 0.817945 | 0.860341 | 0.808509 | 0.669751 |
| CNN | 0.769000 | 0.770023 | 0.692967 | 0.770023 | 0.615506 | 0.770427 | 0.39065 |
| MST-m6A (T5) | 0.599033 | 0.590455 | 0.559427 | 0.591549 | 0.38399 | 0.734864 | 0.126923 |
| deepSRAMP (seqonly) | 0.563431 | 0.541754 | 0.546505 | 0.556966 | 0.454683 | 0.638327 | 0.094619 |
| CLSM6A (T9) | 0.745374 | 0.609362 | 0.633576 | 0.669809 | 0.526888 | 0.740263 | 0.273449 |

**Table S4. DeLong tests comparing M6AFormer with the comparator models**

| Comparison | AUROC difference | Z statistic | P value |
|---|---|---|---|
| M6AFormer vs CLSM6A | +0.1712 | 47.18 | $< 2.2 \times 10^{-16}$ |
| M6AFormer vs MST-m6A | +0.3175 | 89.44 | $< 2.2 \times 10^{-16}$ |
| M6AFormer vs deepSRAMP | +0.3532 | 96.99 | $< 2.2 \times 10^{-16}$ |

**Table S5. Reproduced performance of each comparator on its original held-out test set.**

| Tool | Variant | Metric | Reproduced | Published |
|---|---|---|---|---|
| deepSRAMP | seqonly | AUROC | 0.9586 | 0.967 |
| deepSRAMP | seqonly | AUPRC | 0.6996 | 0.746 |
| MST-m6A | A549 | Accuracy | 0.7834 | within 0.6098-0.8855 |
| MST-m6A | CD8T | Accuracy | 0.7746 | within 0.6098-0.8855 |
| MST-m6A | HCT116 | Accuracy | 0.6098 | within 0.6098-0.8855 |
| MST-m6A | HEK293 | Accuracy | 0.7502 | within 0.6098-0.8855 |
| MST-m6A | HEK293T | Accuracy | 0.7201 | within 0.6098-0.8855 |
| MST-m6A | HeLa | Accuracy | 0.6789 | within 0.6098-0.8855 |
| MST-m6A | HepG2 | Accuracy | 0.6461 | within 0.6098-0.8855 |
| MST-m6A | MOLM13 | Accuracy | 0.814 | within 0.6098-0.8855 |
| MST-m6A | brain | Accuracy | 0.8713 | within 0.6098-0.8855 |
| MST-m6A | kidney | Accuracy | 0.8855 | within 0.6098-0.8855 |
| MST-m6A | liver | Accuracy | 0.8639 | within 0.6098-0.8855 |
| CLSM6A | liver | AUROC | 0.9168 | within 0.6090-0.9168 |
| CLSM6A | kidney | AUROC | 0.9125 | within 0.6090-0.9168 |
| CLSM6A | brain | AUROC | 0.9116 | within 0.6090-0.9168 |
| CLSM6A | MOLM13 | AUROC | 0.8674 | within 0.6090-0.9168 |
| CLSM6A | A549 | AUROC | 0.8451 | within 0.6090-0.9168 |
| CLSM6A | CD8T | AUROC | 0.8431 | within 0.6090-0.9168 |
| CLSM6A | HEK293 | AUROC | 0.8068 | within 0.6090-0.9168 |
| CLSM6A | HEK293T | AUROC | 0.7654 | within 0.6090-0.9168 |
| CLSM6A | HeLa | AUROC | 0.7271 | within 0.6090-0.9168 |
| CLSM6A | HepG2 | AUROC | 0.6466 | within 0.6090-0.9168 |
| CLSM6A | HCT116 | AUROC | 0.609 | within 0.6090-0.9168 |

**Table S6. Cell line–matched evaluation of M6AFormer and comparator models in four cell lines**

| Cell line | Model | AUROC | AUPRC | Acc | Precision | Sen | Spe | MCC |
|---|---|---|---|---|---|---|---|---|
| A549 | M6Aformer | 0.9525 | 0.9471 | 0.864 | 0.8053 | 0.9601 | 0.7679 | 0.7418 |
| A549 | CNN | 0.8142 | 0.8022 | 0.7258 | 0.7606 | 0.6591 | 0.7925 | 0.4558 |
| A549 | CLSM6A(A549) | 0.5712 | 0.5297 | 0.5013 | 0.5033 | 0.2017 | 0.8009 | 0.0033 |
| A549 | MST-m6A(A549) | 0.6684 | 0.6297 | 0.609 | 0.6466 | 0.4806 | 0.7374 | 0.2255 |
| HEK293T | M6AFormer | 0.9497 | 0.9511 | 0.8593 | 0.8104 | 0.938 | 0.7806 | 0.7276 |
| HEK293T | CNN | 0.7617 | 0.7639 | 0.6932 | 0.6904 | 0.7004 | 0.6859 | 0.3863 |
| HEK293T | CLSM6A(HEK293T) | 0.6396 | 0.537 | 0.504 | 0.5087 | 0.2352 | 0.7728 | 0.0095 |
| HEK293T | MST-m6A(HEK293T) | 0.5928 | 0.5716 | 0.5518 | 0.582 | 0.3678 | 0.7358 | 0.1115 |
| HeLa | M6Aformer | 0.9376 | 0.9232 | 0.8613 | 0.8056 | 0.9525 | 0.7702 | 0.735 |
| HeLa | CNN | 0.7865 | 0.7686 | 0.7115 | 0.68 | 0.7989 | 0.624 | 0.4295 |
| HeLa | CLSM6A(HeLa) | 0.4816 | 0.5121 | 0.5059 | 0.5113 | 0.2654 | 0.7463 | 0.0134 |
| HeLa | MST-m6A(HeLa) | 0.6101 | 0.6048 | 0.5715 | 0.619 | 0.3717 | 0.7712 | 0.1559 |
| HepG2 | M6AFormer | 0.9494 | 0.9484 | 0.8596 | 0.806 | 0.9472 | 0.7721 | 0.7306 |
| HepG2 | CNN | 0.8162 | 0.7996 | 0.7485 | 0.7287 | 0.7917 | 0.7053 | 0.4988 |
| HepG2 | CLSM6A(HepG2) | 0.6414 | 0.6446 | 0.6042 | 0.6113 | 0.5722 | 0.6362 | 0.2087 |
| HepG2 | MST-m6A(HepG2) | 0.6461 | 0.6013 | 0.6098 | 0.6249 | 0.5493 | 0.6702 | 0.2211 |

**Table S7. Ablation study of M6AFormer architectural components**

| Model | AUROC | AUPRC | Acc | Precision | Sen | Spe | MCC |
|---|---|---|---|---|---|---|---|
| Full model | 0.9163 | 0.9156 | 0.8286 | 0.7929 | 0.8895 | 0.7677 | 0.6621 |
| w/o Transformer encoder | 0.8573 | 0.8319 | 0.7742 | 0.7202 | 0.8968 | 0.6516 | 0.5656 |
| w/o Residual Conv | 0.9039 | 0.9043 | 0.8151 | 0.7805 | 0.8767 | 0.7534 | 0.6350 |
| w/o Multi-scale Conv | 0.9102 | 0.9098 | 0.8210 | 0.7835 | 0.8869 | 0.7550 | 0.6476 |
| w/o Attention Pool | 0.9100 | 0.9100 | 0.8251 | 0.7981 | 0.8702 | 0.7799 | 0.6528 |

**Table S8. S-LDSC heritability enrichment of predicted m$^6$A sites across traits**

| Trait | Annotation | Prop. SNPs | Coefficient | Coefficient SE | Enrichment | Enrichment SE | Enrichment p |
|---|---|---|---|---|---|---|---|
| AD | Annotated | 0.0379 | 9.82E-09 | 1.02E-08 | 6.47 | 1.61 | 0.000681 |
| AD | Unannotated | 0.01 | 6.3E-09 | 2.88E-08 | 6.45 | 5.51 | 0.33 |
| SCZ | Annotated | 0.0379 | 8.22E-09 | 4.45E-08 | 2.23 | 0.58 | 0.034 |
| SCZ | Unannotated | 0.01 | 1.57E-07 | 7.72E-08 | 3.72 | 1.04 | 0.01 |
| T2D | Annotated | 0.0379 | -7.09E-09 | 1.57E-08 | 1.9 | 1.26 | 0.48 |
| T2D | Unannotated | 0.01 | 1.53E-09 | 2.32E-08 | 1.97 | 1.87 | 0.604 |
| IBD | Annotated | 0.0379 | 7.16E-08 | 1.09E-07 | 4.91 | 1.48 | 0.007 |
| IBD | Unannotated | 0.01 | 6.32E-08 | 2.26E-07 | 4.46 | 3.41 | 0.31 |
| RA | Annotated | 0.0379 | 1.25E-08 | 4.76E-08 | 3.29 | 1.15 | 0.05 |
| RA | Unannotated | 0.01 | -5.28E-08 | 8.8E-08 | 0.66 | 3.05 | 0.911 |
| Height | Annotated | 0.0379 | 1.26E-07 | 9.2E-08 | 4.93 | 0.67 | 3.38E-08 |
| Height | Unannotated | 0.01 | 3.01E-07 | 3.56E-07 | 6.76 | 3.84 | 0.136 |
| BC | Annotated | 0.0379 | 2.02E-08 | 3.15E-08 | 3.35 | 1.17 | 0.048 |
| BC | Unannotated | 0.01 | 8.49E-09 | 5.69E-08 | 2.49 | 2.58 | 0.566 |
| LC | Annotated | 0.0379 | 4.7E-09 | 3.31E-08 | 6.22 | 1.65 | 0.002 |
| LC | Unannotated | 0.01 | 1.48E-07 | 1.01E-07 | 15.06 | 6.92 | 0.055 |
| MDD | Annotated | 0.0379 | 1.26E-08 | 7.59E-09 | 2.43 | 0.47 | 0.002 |
| MDD | Unannotated | 0.01 | 6.4E-09 | 1.52E-08 | 2.06 | 1.19 | 0.371 |
| HCC | Annotated | 0.0379 | -7.87E-10 | 2.2E-09 | 8.83 | 4.97 | 0.027 |
| HCC | Unannotated | 0.01 | 7.92E-09 | 7.51E-09 | 29.7 | 18.39 | 0.203 |

**Table S9. GARFIELD enrichment of predicted $m^6A$ sites across GWAS traits**

| Trait | Annotation | Fold Enrichment | Empirical p | BH-adjusted p | N annot. at threshold | N annot. total | N threshold SNPs |
|---|---|---|---|---|---|---|---|
| AD | Annotated | 4.4532 | <1E-05 | 1.22E-05 | 141 | 49826 | 406 |
| AD | Unannotated | 9.211 | <1E-05 | 1.22E-05 | 26 | 4442 | 406 |
| BC | Annotated | 2.7557 | <1E-05 | 1.22E-05 | 219 | 37967 | 1057 |
| BC | Unannotated | 4.8001 | 2.00E-05 | 2.31E-05 | 36 | 3583 | 1057 |
| Height | Annotated | 2.2794 | <1E-05 | 1.22E-05 | 2930 | 17422 | 9327 |
| Height | Unannotated | 2.3659 | <1E-05 | 1.22E-05 | 387 | 2217 | 9327 |
| IBD | Annotated | 3.9807 | <1E-05 | 1.22E-05 | 229 | 42983 | 734 |
| IBD | Unannotated | 7.9185 | <1E-05 | 1.22E-05 | 42 | 3963 | 734 |
| LC | Annotated | 4.1115 | <1E-05 | 1.22E-05 | 52 | 29902 | 157 |
| LC | Unannotated | 10.6647 | 4.00E-05 | 4.29E-05 | 13 | 2882 | 157 |
| HCC | Annotated | 4.5932 | <1E-05 | <1E-05 | 25 | 35976 | 68 |
| HCC | Unannotated | 3.8318 | 0.346535 | 3.47E-01 | 2 | 3450 | 68 |
| MDD | Annotated | 2.3308 | <1E-05 | 1.22E-05 | 136 | 23519 | 614 |
| MDD | Unannotated | 3.3417 | 1.61E-02 | 1.65E-02 | 21 | 2533 | 614 |
| RA | Annotated | 3.0835 | <1E-05 | 1.22E-05 | 187 | 41797 | 809 |
| RA | Unannotated | 7.3396 | <1E-05 | 1.22E-05 | 41 | 3850 | 809 |
| SCZ | Annotated | 3.3567 | <1E-05 | 1.22E-05 | 281 | 28942 | 1035 |
| SCZ | Unannotated | 6.5888 | <1E-05 | 1.22E-05 | 53 | 2781 | 1035 |
| T2D | Annotated | 2.4418 | <1E-05 | 1.22E-05 | 165 | 13057 | 477 |
| T2D | Unannotated | 2.6806 | 3.10E-02 | 3.10E-02 | 24 | 1730 | 477 |

**Table S10. Composition of the training, validation, and benchmark datasets for M6AFormer**

| Partition | Positive | Negative (DRACH) | Negative (random) | Total (DRACH config) | Total (random config) |
|---|---|---|---|---|---|
| Training | 659,850 | 659,850 | 659,850 | 1,319,700 | 1,319,700 |
| Benchmark | 34,728 | 34,728 | 34,728 | 69,546 | 69,546 |
| Total | 694,578 | 694,578 | 694,578 | 1,389,156 | 1,389,156 |

**Table S11. M6AFormer model architecture and training hyperparameters**

| Parameter | Value | Type |
|---|---|---|
| Input encoding | One-hot (A, C, G, U/T) | Architecture |
| Window lengths evaluated | 201nt, 801nt | |
| Latent dimension (d_model) | 128 | |
| Attention heads (nhead) | 4 | |
| Transformer encoder layers | 2 | |
| Dropout | 0.1 | |
| Pooling | Attention + mean + max (concatenated) | |
| Output | Sigmoid (single logit, BCEWithLogitsLoss) | |
| Framework | PyTorch 2.12.1 | Training |
| Optimizer | AdamW | |
| Initial learning rate | $1 \times 10^{-4}$ | |
| Weight decay | $1 \times 10^{-3}$ | |
| Batch size | 64 | |
| Gradient accumulation steps | 1 | |
| Scheduler | Warm-up cosine | |
| Warm-up ratio | 0.1 | |
| Minimum LR ratio | 0.001 | |
| Max epochs | 50 | |
| Patience | 8 | |
| Positive class weight | 1.0 | |
| Inner validation ratio | 0.1 | |
| Threshold selection metric | F1 | |
| Bootstrap resamples (CI) | 1000 | |

**Table S12. GWAS summary statistics used for S-LDSC and GARFIELD analyses**

| Trait | Accession | Source | Sample size | $h^2$ (SE) | $h^2$ Z-score |
|---|---|---|---|---|---|
| AD | GCST90027158 | GWAS Catalog | 487,511 | 0.0261(0.0027) | 9.67 |
| SCZ | PGC3 wave3 EUR | PGC | 130,644 | 0.3702(0.0133) | 27.83 |
| T2D | GCST006867 | GWAS Catalog | 566,219 | 0.0586(0.0044) | 13.32 |
| IBD | GCST004131 | GWAS Catalog | 59,957 | 0.3236(0.0242) | 13.37 |
| RA | GCST90132223 | GWAS Catalog | 97,173 | 0.1457(0.0120) | 12.14 |
| Height | GCST006901 | GWAS Catalog | 693,529 | 0.5277(0.0214) | 24.66 |
| BC | GCST010098 | GWAS Catalog | 247,173 | 0.1116(0.0107) | 10.43 |
| LC | GCST004748 | GWAS Catalog | 85,716 | 0.0787(0.0110) | 7.15 |
| MDD | GCST90726344 | GWAS Catalog | 514,455 | 0.0656(0.0029) | 22.62 |
| HCC | GCST90809296 | GWAS Catalog | 1,865,284 | 0.0019(0.0009) | 2.11 |

**Table S13. Synthetic nucleotide sequences used in this study**

| Name | Sequence (5'-3') | Assay |
|---|---|---|
| ARHGAP15_val_UP | tagccagtaccgtagtgcgtgTGAACGTATTCAGTAGCTTG | SELECT |
| ARHGAP15_val_DOWN | CTTGTCTGTCAGTCTTCCTCcagaggctgagtcgctgcat | |
| ARHGAP15_neg_UP | tagccagtaccgtagtgcgtgAATATGATGAAGTCAATATC | |
| ARHGAP15_neg_DOWN | GACTGTAGAAGGAACTCATTcagaggctgagtcgctgcat | |
| SERPINB10_val_UP | tagccagtaccgtagtgcgtgTTTTTCACACTGTAAGATGG | |
| SERPINB10_val_DOWN | CTTGTTTGTTGAGAGATATGcagaggctgagtcgctgcat | |
| SERPINB10_neg_UP | tagccagtaccgtagtgcgtgCATTGTGAAATGCATACGTT | |
| SERPINB10_neg_DOWN | TCTCTCCATATATCGCATTGcagaggctgagtcgctgcat | |
| BPI_val_UP | tagccagtaccgtagtgcgtgATGAATAATGTTTCCTCGTG | |
| BPI_val_DOWN | TTAGATCATTTCTGAATCTGcagaggctgagtcgctgcat | |
| BPI_neg_UP | tagccagtaccgtagtgcgtgCACAGAATCTATTTTGGTCA | |
| BPI_neg_DOWN | TACTAGAGAGAAAGAAAATGcagaggctgagtcgctgcat | |
| TARP_val_UP | tagccagtaccgtagtgcgtgCGATACATCTGTGTTCTTTG | |
| TARP_val_DOWN | CCAGTGACTTTTCTGGCACCcagaggctgagtcgctgcat | |
| TARP_neg_UP | tagccagtaccgtagtgcgtgGGCGTGTAAAATGAAAGGCT | |
| TARP_neg_DOWN | CCAGGCAGTTATCTGATTAAcagaggctgagtcgctgcat | |
| PKLR_val_UP | tagccagtaccgtagtgcgtgGGGCTAGATGACAGTTATAG | |
| PKLR_val_DOWN | CTCAGATAGGCCTCAGGTAGcagaggctgagtcgctgcat | |
| PKLR_neg_UP | tagccagtaccgtagtgcgtgTAAAGCAAGGGGAAGACTCC | |
| PKLR_neg_DOWN | CGGCATAAGTGGACCTGGCGcagaggctgagtcgctgcat | |
| MMP11_val_UP | tagccagtaccgtagtgcgtgGCCACAAAGATGGCCATGGG | |
| MMP11_val_DOWN | CTCTAGCCTGATATTCGTGGcagaggctgagtcgctgcat | |
| MMP11_neg_UP | tagccagtaccgtagtgcgtgCCCCATTTGACTGTGAACTT | |
| MMP11_neg_DOWN | TTGGCCTGGATTCTGGAGAAcagaggctgagtcgctgcat | |
| NEU4_val_UP | tagccagtaccgtagtgcgtgGAGCTCTGAGGAGCAACTTG | |
| NEU4_val_DOWN | TTCTAATCCACAGGATCCTGcagaggctgagtcgctgcat | |
| NEU4_neg_UP | tagccagtaccgtagtgcgtgACACAAGCACGATTCCTTTA | |
| NEU4_neg_DOWN | TTCGAGCCAGAAAGCAGCCAcagaggctgagtcgctgcat | |
| PFDN6_val_UP | tagccagtaccgtagtgcgtgTCAGGAAACAGAGCCAGGTG | |
| PFDN6_val_DOWN | TTTTACATTTTATTAGCTACcagaggctgagtcgctgcat | |
| PFDN6_neg_UP | tagccagtaccgtagtgcgtgAACTCACATTTCAGCTGTGA | |
| PFDN6_neg_DOWN | ATAGTCCAGCCTCTTCCCTAcagaggctgagtcgctgcat | |
| qPCRF | ATGCAGCGACTCAGCCTCTG | |
| qPCRR | TAGCCAGTACCGTAGTGCGTG | |
| NEU4_F | ACCGCCTGGTGCTGAGGAGG | NEU4 mRNA expression |
| NEU4_R | TGAAGAAGAGGAAGACGGTGCC | |
| GAPDH_F | ACCCAGAAGACTGTGGATGG | |
| GAPDH_R | TTCAGCTCAGGGATGACCTT | |

| | | |
|---|---|---|
| NEU4 gRNA | GCAGAGATCCAGTTTGGTTAATTAAGGTACCGAG<br>GGCCTATTTCCCATGATTCCTTCATATTTGCATATA<br>CGATACAAGGCTGTTAGAGAGATAATTAGAATTA<br>ATTTGACTGTAAACACAAAGATATTAGTACAAAA<br>TACGTGACGTAGAAAGTAATAATTTCTTGGGTAG<br>TTTGCAGTTTTAAAATTATGTTTTAAAATGGACTA<br>TCATATGCTTACCGTAACTTGAAAGTATTTCGATT<br>TCTTGGCTTTATATATCTTGTGGAAAGGACGAAA<br>AAACACCGAACCCCTACCAACTGGTCGGGGTTT<br>GAAACgGAGCTCTGAGGAGCAACTTGTTTCTAAT<br>CCTTTTTTGCTAGCTAGGTCTTGAAAGGAGTGGG<br>AATT [a] | Site-specific m$^6$A removal |
| Non-targeting gRNA | GCAGAGATCCAGTTTGGTTAATTAAGGTACCGAG<br>GGCCTATTTCCCATGATTCCTTCATATTTGCATATA<br>CGATACAAGGCTGTTAGAGAGATAATTAGAATTA<br>ATTTGACTGTAAACACAAAGATATTAGTACAAAA<br>TACGTGACGTAGAAAGTAATAATTTCTTGGGTAG<br>TTTGCAGTTTTAAAATTATGTTTTAAAATGGACTA<br>TCATATGCTTACCGTAACTTGAAAGTATTTCGATT<br>TCTTGGCTTTATATATCTTGTGGAAAGGACGAAA<br>AAACACCGAACCCCTACCAACTGGTCGGGGTTT<br>GAAACgCGTCTGGCCTTCCTGTAGCCAGCTTTCA<br>TCTTTTTTGCTAGCTAGGTCTTGAAAGGAGTGGG<br>AATT [a] | |

[a] Colours indicate the structural parts of the gRNA construct. Grey, 30-bp buffer, U6 promoter and DR30 direct repeat. Red, gRNA spacer, where the lowercase g marks the additional 5' guanine required for U6 transcription. Blue, poly-T terminator and 30-bp buffer. For SELECT oligonucleotides, lowercase bases denote the shared adaptor sequences and uppercase bases the target-specific region.
